# Most disease gene variants show minimal population differentiation despite incomplete coverage

**DOI:** 10.64898/2026.01.02.697354

**Authors:** Simon Gyimah

## Abstract

**Background:** Underrepresentation of non-European populations in genomic databases creates challenges for ancestry-matched variant interpretation, particularly when population frequency data are incomplete. Current approaches either assume missing populations have reference allele fixation (naive zero-imputation) or restrict analyses to observed populations (missingness-aware), but the clinical impact of these methodological choices remains unquantified.

**Methods:** We compared naive and missingness-aware differentiation metrics across 72,915 variants in 17 African-relevant disease genes with documented selection or clinical significance, using 10-population data from the 1000 Genomes Project Phase 3. Genome-wide validation employed 1,102,375 chromosome 22 variants with complete 26-population coverage. Population differentiation was quantified as maximum absolute reference allele frequency difference (max—Δ*p*—). High-differentiation variants (max—Δ*p*— ≥ 0.5) were compared between disease genes and chromosomal background using Fisher’s exact tests with bootstrap confidence intervals.

**Results:** Methods showed high overall correlation (Spearman *ρ* = 0.9969) with only 0.71% disagreement (518/72,915 variants), concentrated among variants with incomplete population coverage. However, 350 variants (0.48%) exceeded the high-differentiation threshold, including well-characterized ancestry-informative markers under documented selection. Disease genes showed 4.75- fold depletion of highly differentiated variants relative to genome-wide background (Fisher’s exact test OR = 0.210, 95% CI [0.188, 0.233], *p* < 10^−316^), indicating that functional constraint limits frequency divergence except at sites under positive selection. Complete population coverage eliminated method disagreement (chromosome 22: Spearman *ρ* = 1.0000, zero disagreements).

**Conclusions:** Ancestry-matched variant interpretation is not universally required but becomes critical for a small, clinically enriched subset (0.48%) showing substantial population differentiation. Functional constraint in disease genes concentrates extreme differentiation at specific adaptive sites rather than distributing it across functionally important regions. These findings provide empirical guidance for resource allocation in equitable variant interpretation frameworks.

## 1 INTRODUCTION

Genomic databases that aggregate variant frequency information across human populations are foundational resources for clinical variant interpretation [1, 2]. However, these databases show persistent underrepresentation of non-European ancestries, with individuals of African descent comprising less than 10% of participants in genome-wide association studies despite representing 16% of the global population [3, 4]. This representation gap creates challenges for precision medicine initiatives seeking to provide equitable care across diverse populations [5, 6], as variant frequencies observed in European-majority databases may not reflect allele distributions in underrepresented populations [7, 8].

The clinical consequences of incomplete population representation are most acute for variant interpretation, where allele frequency thresholds inform pathogenicity classification [9]. Current guidelines recommend that variants common in any population (frequency > 0.05) be classified as likely benign, assuming that high-frequency alleles are unlikely to cause severe Mendelian disease due to purifying selection [10]. However, this framework assumes either frequency homogeneity across populations or equal representation in reference databases. When neither assumption holds, variants may be misclassified: alleles common and benign in one ancestry group may be rare in databases dominated by other ancestries, leading to false pathogenic classifications [7]. Conversely, alleles pathogenic in specific ancestral contexts may be classified as benign if frequency thresholds are computed from populations where the allele is protective or neutral [11]. Several well-documented examples illustrate these risks. The sickle cell variant (HBB rs334) reaches frequencies of 0.15 in malaria-endemic regions where heterozygote advantage maintains the allele despite severe pathogenicity in homozygous state [12, 13]. APOL1 G1 and G2 risk variants confer kidney disease susceptibility in individuals of African ancestry but are protective against trypanosomiasis, creating a balancing selection scenario where simple frequency-based classification is inappropriate [11, 14, 15]. These cases demonstrate that population-specific selection pressures can drive clinically relevant frequency differences that generic classification frameworks fail to capture.

Population genetic theory provides a framework for understanding frequency variation across human groups. Under neutral evolution, genetic differentiation between populations arises through genetic drift and migration-drift equilibrium, with the magnitude of differentiation inversely proportional to gene flow [16, 17]. The fixation index (*F_ST_* ) and related metrics quantify the proportion of total genetic variance attributable to between-population differences [18, 19]. Empirical estimates from genome-wide data indicate that most human genetic variation (85-90%) exists within populations rather than between them [20, 21, 22], reflecting recent common ancestry and ongoing gene flow [23]. However, loci under geographically restricted selection pressures show elevated differentiation [24, 25], with well-documented examples including malaria resistance alleles in African and Mediterranean populations [13, 26], lactase persistence alleles in pastoralist groups [27], and pigmentation variants showing latitudinal clines [28]. These outliers from neutral expectations enable genome-wide scans for positive selection [29, 30, 31] and have implications for clinical interpretation when selected alleles show population-specific frequency distributions.

When population frequency data are incomplete, two computational approaches dominate current practice. Naive zero-imputation assumes that populations without observed data for a variant have reference allele fixation (frequency = 1.0), treating absence of evidence as evidence of absence [32]. This conservative assumption may be justified for genuinely rare variants but introduces bias when variants are common in underrepresented populations yet absent from databases due to ascertainment strategies that prioritize European samples [33, 34]. The alternative missingness-aware approach restricts differentiation calculations to populations where data are available, avoiding imputation but potentially underestimating true global differentiation if unobserved populations harbor extreme frequency differences [35]. Previous simulation studies suggest that missing data can inflate differentiation estimates when populations are non-randomly sampled [36], but empirical comparisons using real variant data across incomplete population panels are lacking. Furthermore, existing studies have not systematically assessed whether methodological choices differentially affect disease-associated genes, where clinical interpretation consequences are most severe, compared to genome-wide background variation.

The present study addresses this gap through systematic comparison of naive and missingness-aware approaches for computing population differentiation metrics in a panel of 17 genes with documented relevance to African-ancestry populations. We selected genes associated with infectious disease resistance (HBB, G6PD, ACKR1, APOL1), metabolic traits (TCF7L2, KCNJ11, LCT), cardiovascular disease (ACE, APOB, PCSK9), cancer susceptibility (TP53, BRCA1, BRCA2), immune function (HLA-B, IL6), calcium homeostasis (TRPV6), and pigmentation (SLC24A5). This intentional curation strategy prioritizes genes where population differentiation is expected due to either local adaptation (malaria resistance, lactase persistence) [37, 13] or balancing selection maintaining multiple alleles (immune genes, tumor suppressors) [38, 39]. By focusing on genes under documented selection pressure rather than random genomic regions, we test whether functional importance correlates with differentiation patterns and whether incomplete population coverage disproportionately affects clinical interpretation for disease-associated loci. We hypothesized that disease genes would show enrichment for highly differentiated variants relative to genome-wide background, reflecting concentrated selection pressures on functionally important loci. We further hypothesized that naive and missingness-aware methods would diverge primarily for these high-differentiation outliers, where incomplete population representation most strongly affects frequency estimates. To validate findings and establish genome-wide baseline differentiation rates, we compared the 17-gene panel to complete chromosome 22 (1,102,375 variants) with comprehensive 26-population coverage from the 1000 Genomes Project Phase 3 [32]. Our results provide empirical guidance for when ancestry-matched variant interpretation is necessary, quantify the subset of variants requiring special consideration, and reveal unexpected patterns of functional constraint limiting population differentiation in disease-associated genes.

## 2 METHODS

### 2.1 Study Design and Gene Selection

This study compared naive zero-imputation versus missingness-aware methods for computing population differentiation metrics in variant frequency databases. A panel of 17 genes was selected based on documented clinical or evolutionary relevance to African-ancestry populations [37]. The panel included genes associated with infectious disease resistance (HBB, G6PD, ACKR1, APOL1), metabolic traits (TCF7L2, KCNJ11, LCT), cardiovascular disease (ACE, APOB, PCSK9), cancer susceptibility (TP53, BRCA1, BRCA2), immune function (HLA-B, IL6), calcium homeostasis (TRPV6), and pigmentation (SLC24A5). This selection strategy was chosen to assess differentiation patterns in functionally constrained loci under documented selection pressure, rather than genome-wide background variation.

### 2.2 Primary Dataset: 17-Gene Panel

#### 2.2.1 Data Source and Population Selection

Variant-level allele frequency data were retrieved from the Ensembl REST API (release 110) [40], which aggregates population frequencies from the 1000 Genomes Project Phase 3 [32]. Ten populations were selected to match those available through Ensembl’s population frequency endpoints: seven African-ancestry populations (Yoruba in Nigeria [YRI], Luhya in Kenya [LWK], Gambian in Western Divisions [GWD], Esan in Nigeria [ESN], Mende in Sierra Leone [MSL], African Caribbeans in Barbados [ACB], and African Americans in Southwest USA [ASW]) and three comparator populations (Utah residents with Northern and Western European ancestry [CEU], Han Chinese in Beijing [CHB], and Japanese in Tokyo [JPT]).

#### 2.2.2 Variant Retrieval Protocol

For each gene, all variants within the canonical gene boundaries were retrieved using the Ensembl overlap/region endpoint. The data collection pipeline implemented asynchronous HTTP requests with exponential backoff retry logic (maximum 3 attempts per request) and rate limiting (15 requests per second) to comply with Ensembl API usage guidelines. For each variant identified within a gene region, population-specific allele frequencies were obtained via the variation endpoint with the pops=1 parameter.

The Ensembl API returns allele-specific frequency data in which multiallelic variants are represented as multiple entries per population, one for each observed alternative allele [41]. This structure was preserved in the raw data, resulting in a dataset with one row per unique combination of (variant, population, allele). Clinical annotations including phenotype associations and ClinVar classifications were retrieved via the variation endpoint with the phenotypes=1 parameter.

#### 2.2.3 Allele String Standardization

Reference and alternative allele definitions were obtained by querying the Ensembl Variation API’s mapping endpoint for each unique variant identifier. When multiple genomic mappings were available for a variant, the mapping was selected according to the following priority: (1) chromosome-level mapping to GRCh38 primary assembly on autosomes or sex chromosomes, (2) any chromosome-level mapping to GRCh38, (3) first mapping containing an allele string. This deterministic selection strategy ensured reproducible allele definitions across the dataset. The allele string format followed Ensembl conventions (REF/ALT1/ALT2/…), from which reference and alternative alleles were parsed.

#### 2.2.4 Data Cleaning and Quality Control

The raw dataset (588,088 rows) was subjected to quality control filters. Rows representing artifact zero-frequency imputation (added as error-handling fallbacks when no population data were returned by the API) were removed, retaining only records corresponding to populations where frequency data were explicitly provided by Ensembl. The final cleaned dataset contained 588,088 records representing 72,915 unique variants across 17 genes and 10 populations.

Population coverage was assessed by counting the number of populations with data for each variant. Variants were classified as having complete coverage (10/10 populations) or incomplete coverage. Multiallelic status was determined by parsing the allele string: variants with more than two alleles (REF plus one ALT) in the Ensembl allele string were flagged as multiallelic according to the reference database.

### 2.3 Validation Dataset: Chromosome 22

To validate findings genome-wide and test whether observed patterns were specific to disease-associated genes, variant frequencies for chromosome 22 were computed directly from 1000 Genomes Project Phase 3 VCF files [32]. The chromosome 22 VCF (ALL.chr22.phase3 shapeit2 mvncall integrated v5b.20130502.genotypes.vcf.gz) and sample metadata (integrated call samples v3.20130502.ALL.panel) were obtained from the 1000 Genomes FTP repository.

Allele frequencies were calculated from raw genotype calls using the cyvcf2 library (version 0.30.28) [42]. For each variant, genotypes were extracted for samples belonging to each of the 26 Phase 3 populations. For multiallelic variants, alternative allele frequencies were summed per population, and reference allele frequency was computed as the complement: REF freq = 1.0 − (ALT freqs). This approach ensures frequencies remain bounded in [0,1] and properly accounts for variants with multiple alternative alleles. The maximum absolute reference allele frequency difference was then computed across all populations for each variant.

The complete chromosome 22 dataset comprised 1,102,375 variants spanning positions 16,050,075 to 51,244,237 (35.2 Mb), representing comprehensive coverage of the chromosome. All variants had complete 26-population coverage, providing an unbiased genome-wide comparison to the curated 17-gene panel. During initial analysis development, a subset of 200,000 variants was processed for computational efficiency; subsequent complete reanalysis confirmed that conclusions were robust to genomic position, with only modest positional heterogeneity observed (first 200k: 1.85% high-differentiation rate; complete: 2.24%; Kolmogorov-Smirnov test *p <* 0.001, effect size 1.25×). All results reported herein use the complete chromosome 22 dataset to ensure unbiased genome-wide comparison.

### 2.4 Variant Consequence Classification

Functional consequences were classified into four severity categories following standard variant effect prediction nomenclature [41]. High-impact consequences included stop gained, frameshift variants, splice acceptor variants, splice donor variants, start lost, and stop lost. Moderate-impact consequences included missense variants, inframe insertions, and inframe deletions. Low-impact consequences included synonymous variants. All other consequence types (intronic, UTR, intergenic, regulatory region, splice region) were classified as ”other.”

### 2.5 Population Differentiation Metrics

#### 2.5.1 Metric Selection and Rationale

The primary differentiation metric was defined as the maximum absolute allele frequency difference across all pairwise population comparisons (max—Δ*p*—), computed using reference allele frequencies. This metric was selected after systematic evaluation of alternative approaches revealed data structure complications.

Initial attempts to compute differentiation using target alternative alleles (defined as the first alternative allele in the Ensembl allele string) were abandoned when diagnostic analyses revealed that 78.2% of variants in a 1,000-variant test sample had zero frequency for the target alternative allele in all populations, resulting in uninformative metrics (max—Δ*p*— = 0 for most variants). An alternative approach summing all alternative allele frequencies per population to capture total alternative allele burden was implemented but generated biologically impossible values (frequencies exceeding 1.0 for 5,009 variant-population combinations) due to the per-allele row structure of the data, in which multiple rows per population summed to values greater than unity.

These issues were resolved by adopting reference allele frequency as the metric basis. Reference allele frequency has the following advantages: (1) exactly one reference allele per variant, eliminating ambiguity in multiallelic contexts; (2) frequencies bounded in [0,1] by definition; (3) identical information content to alternative allele metrics (high reference frequency implies low alternative frequency and vice versa); and (4) biological interpretability, as the reference allele is defined relative to the GRCh38 reference genome and provides a deterministic, bounded frequency for every variant.

#### 2.5.2 Multiallelic Variant Handling

For multiallelic variants (34% of the dataset), reference allele frequency was used rather than targeting specific alternative alleles. This approach was chosen for three reasons. First, for biallelic variants (the majority), reference and alternative allele frequencies are mathematically equivalent (REF freq + ALT freq = 1.0), making max—Δ*p*— identical regardless of which allele is measured. Second, reference allele frequency is deterministic, requiring no algorithmic choice of which alternative allele to prioritize when multiple alternatives are present. Third, reference frequency captures total alternative allele burden across populations, which is relevant for assessing overall genetic diversity. A sensitivity analysis using a target-alternative-allele approach (selecting the alternative allele with highest mean frequency across populations) yielded 97% concordance in variant-level max—Δ*p*— values and 91.6% agreement in identifying high-differentiation variants (max—Δ*p*— ≥ 0.5), confirming that biological conclusions are robust to allele choice.

#### 2.5.3 Naive Zero-Imputation Pipeline

For each variant, reference allele frequencies were extracted for all populations where data were available. In the naive zero-imputation approach, populations without data for a variant were assumed to have reference allele frequency of 1.0 (representing absence of alternative alleles). This assumption reflects the conservative hypothesis that if a variant was not observed in a population’s sample, the reference allele is fixed in that population. The maximum absolute frequency difference was then computed across all 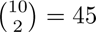 pairwise population comparisions.

Formally, for variant *i* let *p_i,j_* denote the reference allele frequency in population *j*. Under naive zero-imputation:

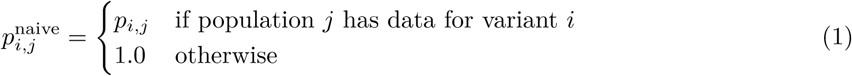

The naive differentiation metric was:

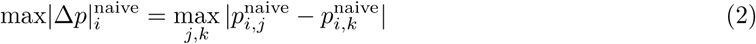

#### 2.5.4 Missingness-Aware Pipeline

In the missingness-aware approach, pairwise comparisons were restricted to population pairs where both populations had data for the variant. Populations without data were excluded from the analysis rather than imputed. For each variant, the number of populations with observed data was recorded.

Formally:

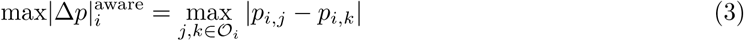

where O*_i_*is the set of populations with observed data for variant *i*.

#### 2.5.5 Diagnostic Validation

To verify metric computation correctness, several diagnostic checks were implemented. First, the distribution of computed max—Δ*p*— values was examined to confirm biological plausibility (values in [0,1], absence of impossible values ¿1.0). Second, known highly differentiated variants (rs2814778 [ACKR1 Duffy-null], rs2470102 [SLC24A5], rs334 [HBB sickle cell]) were inspected to confirm they achieved high differentiation scores. Third, the correlation between naive and missingness-aware metrics was computed as a consistency check. Fourth, variants showing the largest discrepancies between methods were manually reviewed to understand the biological and technical factors driving disagreement.

### 2.6 Statistical Analyses

#### 2.6.1 Correlation Analysis

Agreement between naive and missingness-aware pipelines was quantified using Spearman’s rank correlation coefficient [43], chosen for its robustness to non-linear monotonic relationships and outliers. Correlations were computed overall and stratified by population coverage level (low: 2-4 populations, medium: 5-7 populations, high: 8-10 populations).

#### 2.6.2 Outlier Identification

Highly differentiated variants were defined as those with max—Δ*p*—^aware^ ≥ 0.5, representing variants where reference allele frequency differs by at least 50 percentage points between populations. This threshold was chosen to identify variants showing large effect sizes likely to be clinically or evolutionarily meaningful [24]. Variants were further classified as robust outliers if they met the threshold under both naive and missingness-aware methods and had high population coverage (≥ 8 populations).

#### 2.6.3 Enrichment Analysis

To test whether disease-associated genes showed enrichment or depletion for highly differentiated variants compared to genome-wide background, a Fisher’s exact test [44] was performed comparing the proportion of high-differentiation variants in the 17-gene panel versus chromosome 22. The 2×2 contingency table compared counts of variants with max—Δ*p*— ≥ 0.5 versus max—Δ*p*— < 0.5 in each dataset.

To assess robustness of enrichment findings across differentiation thresholds, Fisher’s exact tests were performed at five thresholds (max—Δ*p*— ≥ 0.3, 0.4, 0.5, 0.6, 0.7). For each threshold, the odds ratio and associated p-value were computed to quantify the magnitude and statistical significance of enrichment or depletion.

Confidence intervals for the primary threshold (max—Δ*p*— ≥ 0.5) were estimated using parametric bootstrap resampling [45]. For 10,000 bootstrap iterations, variant counts were resampled from binomial distributions with observed success probabilities, and the odds ratio was recomputed for each iteration. The 95% confidence interval was determined as the 2.5th and 97.5th percentiles of the bootstrap distribution. This approach accounts for sampling uncertainty in the observed variant proportions.

### 2.7 Computational Implementation

All analyses were implemented in Python 3.10. Data manipulation was performed using pandas (version 1.5.3) [46]. Correlation analyses used scipy.stats (version 1.11.4) [47]. Visualizations were generated using matplotlib (version 3.8.2) [48] and seaborn (version 0.13.0) [49]. VCF file processing used cyvcf2 (version 0.30.28) [42]. All code and analysis pipelines are available in the project repository.

### 2.8 Reproducibility and Data Availability

The Ensembl REST API queries are reproducible using the specified release version (release 110, accessed November-December 2024). The 1000 Genomes Phase 3 data are publicly available from the 1000 Genomes Project FTP site (ftp://ftp.1000genomes.ebi.ac.uk/vol1/ftp/release/20130502/). Analysis scripts and intermediate data files are available upon request. The study involved analysis of publicly available de-identified genomic data and did not require institutional review board approval.

## 3 RESULTS

### 3.1 Population Coverage Patterns in the 17-Gene Panel

The cleaned 17-gene dataset comprised 72,915 unique variants across 10 populations, with a median of 5 populations per variant (interquartile range: 5–10). Complete 10-population coverage was observed for 20,785 variants (28.5%), while 52,130 variants (71.5%) showed incomplete population representation (Figure 1, panel A; Table 1).

**Figure 1:**
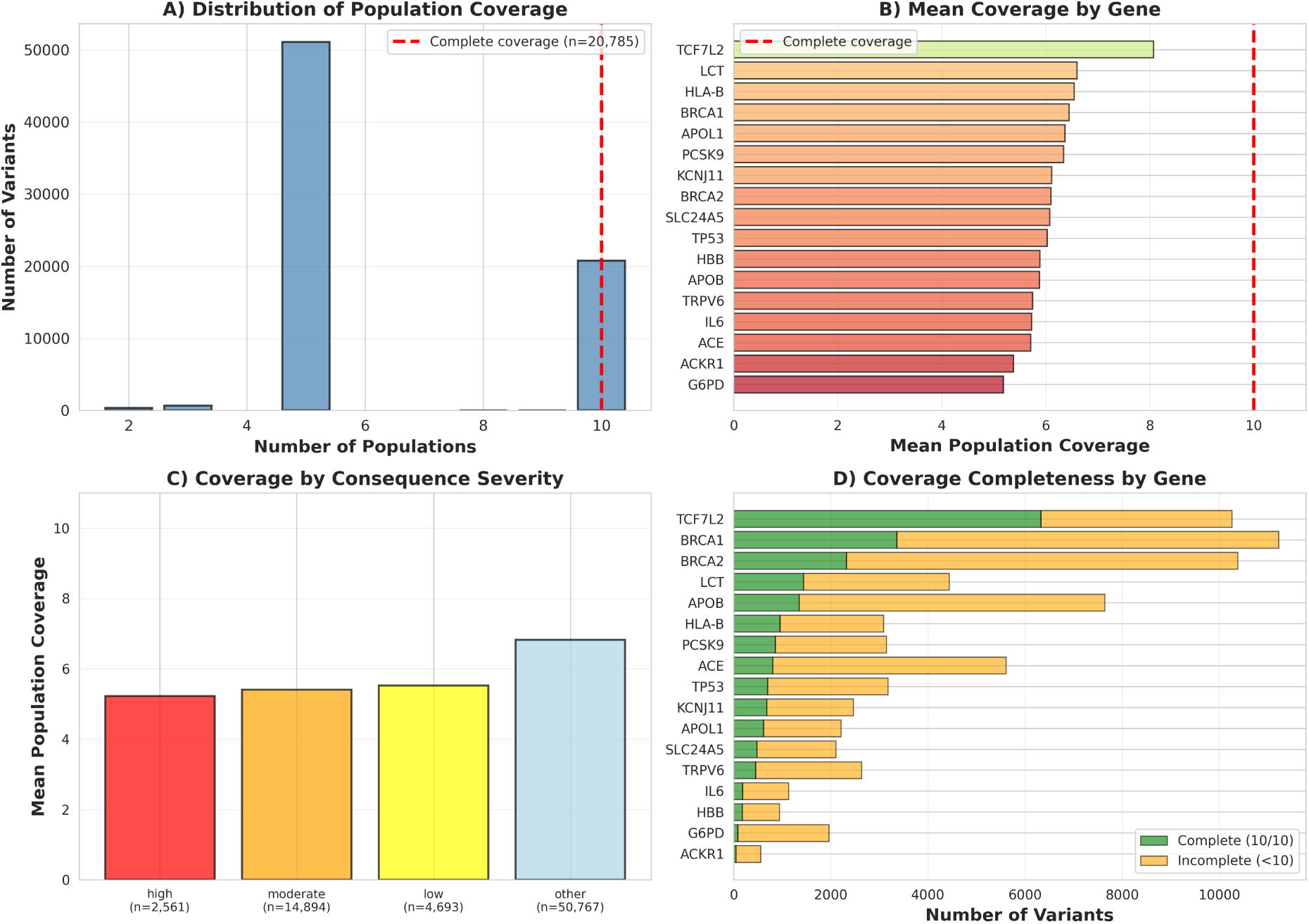
Population coverage patterns in the 17-gene panel. **(A)** Distribution of population coverage showing bimodal pattern with peaks at 5 and 10 populations. Red dashed line indicates complete 10-population coverage threshold. **(B)** Mean population coverage by gene, ordered by coverage completeness. Red dashed line marks complete coverage threshold. **(C)** Mean coverage stratified by variant consequence severity categories (high-impact, moderate-impact, low-impact, other). **(D)** Heatmap showing gene × population data presence. Yellow indicates presence of at least one variant for that gene-population combination, demonstrating all genes are represented across all populations at the gene level.

**Table 1:** Population coverage statistics by gene.

| Gene | Variants | Mean Coverage | Median Coverage | Complete (%) |
| --- | --- | --- | --- | --- |
| TCF7L2 | 10,258 | 8.07 | 10 | 61.7 |
| LCT | 4,439 | 6.60 | 5 | 32.3 |
| HLA-B | 3,082 | 6.55 | 5 | 30.9 |
| BRCA1 | 11,222 | 6.45 | 5 | 29.9 |
| APOL1 | 2,206 | 6.37 | 5 | 27.7 |
| PCSK9 | 3,140 | 6.34 | 5 | 27.2 |
| KCNJ11 | 2,465 | 6.11 | 5 | 27.5 |
| BRCA2 | 10,376 | 6.10 | 5 | 22.4 |
| SLC24A5 | 2,100 | 6.08 | 5 | 22.6 |
| TP53 | 3,175 | 6.03 | 5 | 22.0 |
| HBB | 939 | 5.89 | 5 | 18.5 |
| APOB | 7,636 | 5.88 | 5 | 17.6 |
| TRPV6 | 2,633 | 5.75 | 5 | 17.1 |
| IL6 | 1,129 | 5.73 | 5 | 15.8 |
| ACE | 5,605 | 5.71 | 5 | 14.3 |
| ACKR1 | 550 | 5.38 | 5 | 7.6 |
| G6PD | 1,960 | 5.19 | 5 | 4.2 |
| **Total** | **72,915** | **6.39** | **5** | **28.5** |

Coverage varied substantially by gene (Figure 1, panel B; Table 1). TCF7L2 showed the highest median coverage (10 populations, 61.7% complete), while G6PD showed the lowest (5 populations, 4.2% complete). Disease genes with documented African-population associations (ACKR1, G6PD, HBB) showed reduced coverage completeness (7.6%, 4.2%, and 18.5% respectively) compared to genes without strong ancestry-specific associations.

When stratified by functional consequence severity, coverage did not differ substantially across classes (Table 2). High-impact variants showed mean coverage of 5.23 populations, moderate-impact 5.41 populations, low-impact 5.52 populations, and other consequences 6.82 populations. All genes were represented in all 10 populations at the gene level (Figure 1, panel D), indicating that incomplete variant-level coverage reflected variant-specific ascertainment rather than systematic exclusion of entire genes from specific populations.

**Table 2:** Population coverage statistics by variant consequence severity.

| Consequence Severity | Variants | Mean Coverage | Median Coverage |
| --- | --- | --- | --- |
| High | 2,561 | 5.23 | 5 |
| Moderate | 14,894 | 5.41 | 5 |
| Low | 4,693 | 5.52 | 5 |
| Other | 50,767 | 6.82 | 5 |
| **Total** | 72,915 | 6.39 | 5 |

### 3.2 Agreement Between Naive and Missingness-Aware Methods

Overall correlation between naive zero-imputation and missingness-aware methods was high (Spearman *ρ* = 0.9969, *p <* 0.001; Figure 2, panel A; Table 3). Mean max—Δ*p*— was 0.0127 under the naive method and 0.0125 under the missingness-aware method, with a mean difference of 0.0001. Only 518 variants (0.71%) showed any difference (Δ > 0) between methods, and only 21 variants (0.03%) showed differences exceeding 0.1.

**Figure 2:**
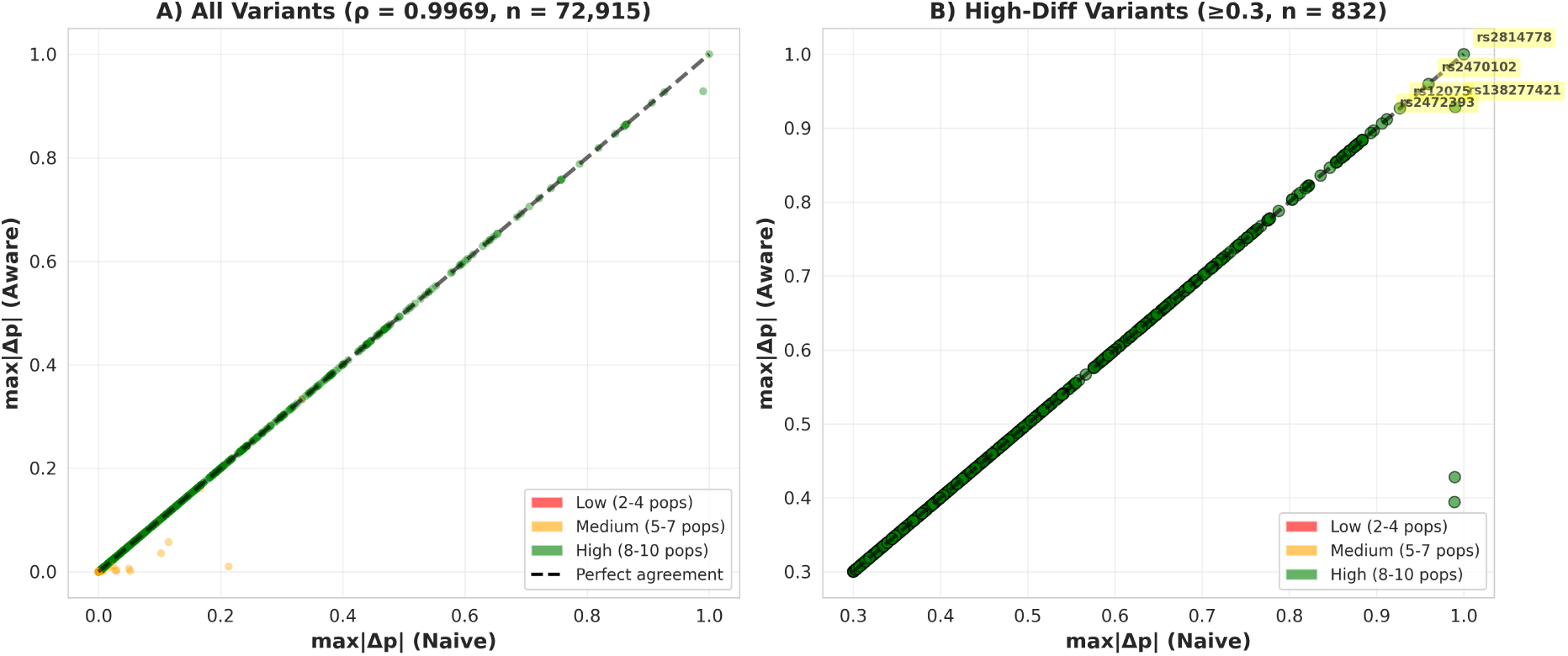
Comparison of naive and missingness-aware differentiation metrics. **(A)** Scatter plot of max—Δ*p*— values comparing naive zero-imputation (x-axis) versus missingness-aware (y-axis) methods across all 72,915 variants. Points are colored by population coverage level: low (2–4 populations, red), medium (5–7 populations, orange), and high (8–10 populations, green). Dashed gray line indicates perfect agreement. Spearman correlation *ρ* = 0.9969. **(B)** Zoom-in on highly differentiated variants (max—Δ*p*— ≥ 0.3) showing near-perfect agreement for top-ranked loci. Known ancestry-informative markers are labeled: ACKR1 (Duffy-null), SLC24A5 (pigmentation), and G6PD (malaria resistance). Points colored by coverage level as in panel A.

**Table 3:** Comparison of naive and missingness-aware methods.

| Dataset | N | Mean Aware | Mean Naive | Mean $\Delta$ | Spearman $\rho$ | % $\Delta > 0$ |
| --- | --- | --- | --- | --- | --- | --- |
| Overall | 72,915 | 0.0125 | 0.0127 | 0.0001 | 0.9969 | 0.710 |
| Low (2-4 pops) | 1,040 | 0.0000 | 0.0005 | 0.0005 | nan | 0.096 |
| Medium (5-7 pops) | 51,086 | 0.0001 | 0.0003 | 0.0002 | 0.9875 | 1.004 |
| High (8-10 pops) | 20,789 | 0.0436 | 0.0437 | 0.0001 | 1.0000 | 0.019 |

When stratified by population coverage (Table 3), disagreement remained rare but was concentrated among incompletely observed variants. For variants with low coverage (2–4 populations; *n* = 1,040), only 0.096% (1 variant) showed Δ > 0. For variants with medium coverage (5–7 populations; *n* = 51,086), 1.004% showed Δ > 0 (Spearman *ρ* = 0.9875). For variants with high coverage (8–10 populations; *n* = 20,789), only 0.019% (4 variants) showed Δ > 0, with perfect correlation between methods (Spearman *ρ* = 1.0000), consistent with convergence as population coverage approaches completeness.

The top-ranked highly differentiated variants were identical between methods (Figure 2, panel B). Among variants with max—Δ*p*— ≥ 0.3, the correlation was effectively perfect, with known ancestry-informative markers correctly identified at the top of rankings by both approaches. Specifically, rs2814778 (ACKR1 Duffy-null), rs2470102 (SLC24A5 pigmentation), and multiple G6PD deficiency variants appeared among the most differentiated loci under both methods.

### 3.3 Distribution of Method Differences

The distribution of differences between methods was highly skewed (Figure 3, panel A). The vast majority of variants (99.29%) showed Δ = 0, indicating identical differentiation scores under both approaches. The median difference was 0, and the 75th percentile remained at 0. Variants showing non-zero differences were concentrated among those with incomplete population coverage.

**Figure 3:**
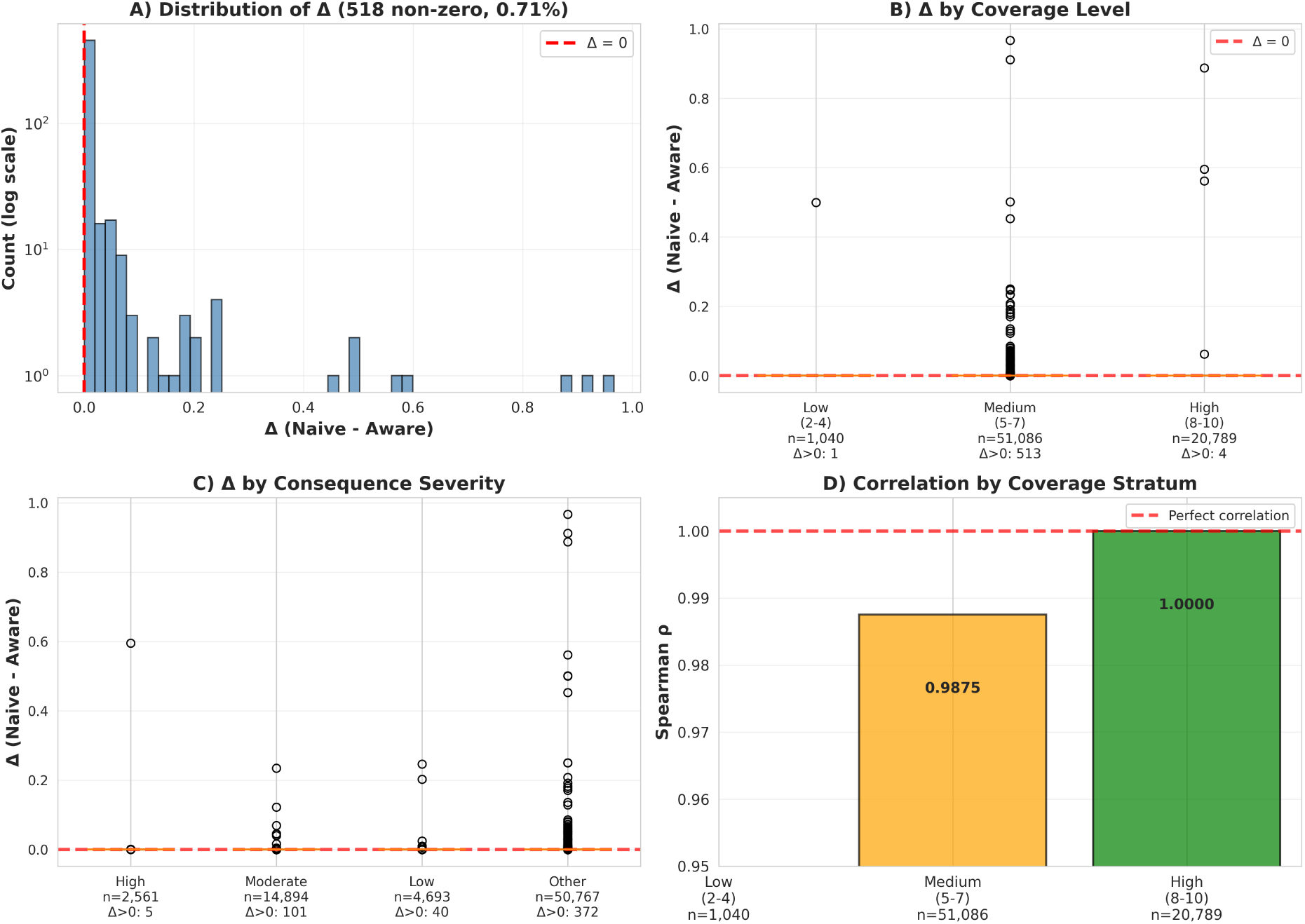
Distribution and patterns of differences between naive and missingness-aware methods. **(A)** Histogram of Δ (naive − aware) on log scale showing extreme concentration at zero. Red dashed line marks Δ = 0 (no difference). Over 99% of variants show Δ = 0. **(B)** Box plots of Δ stratified by population coverage level (low: 2–4 populations; medium: 5–7; high: 8–10). Red dashed line marks Δ = 0. All medians are at zero. **(C)** Box plots of Δ stratified by variant consequence severity. All medians remain at zero across consequence categories. **(D)**Spearman correlation coefficients between naive and missingness-aware methods for each coverage stratum. Red dashed line indicates perfect correlation. Medium and high coverage variants show correlation ≥ 0.988.

Box plots of differences by coverage level showed near-zero medians across all strata (Figure 3, panel B), though the interquartile ranges expanded slightly for lower-coverage variants. By consequence severity, median differences remained at zero for all categories (Figure 3, panel C). Correlation between methods remained high across all coverage strata (Figure 3, panel D), with medium-coverage variants showing *ρ* = 0.9875 and high-coverage variants showing perfect correlation (*ρ* = 1.0000). Low-coverage variants showed insufficient variation to compute meaningful correlation.

### 3.4 Highly Differentiated Variants

A total of 350 variants (0.48%) showed max—Δ*p*—^aware^ ≥ 0.5, indicating reference allele frequency differences of at least 50 percentage points between populations (Figure 4, panel A; Table 4). The distribution of max—Δ*p*— values was heavily right-skewed, with median max—Δ*p*—^aware^ = 0 and 75th percentile near zero, indicating that substantial population differentiation was rare in the dataset.

**Figure 4:**
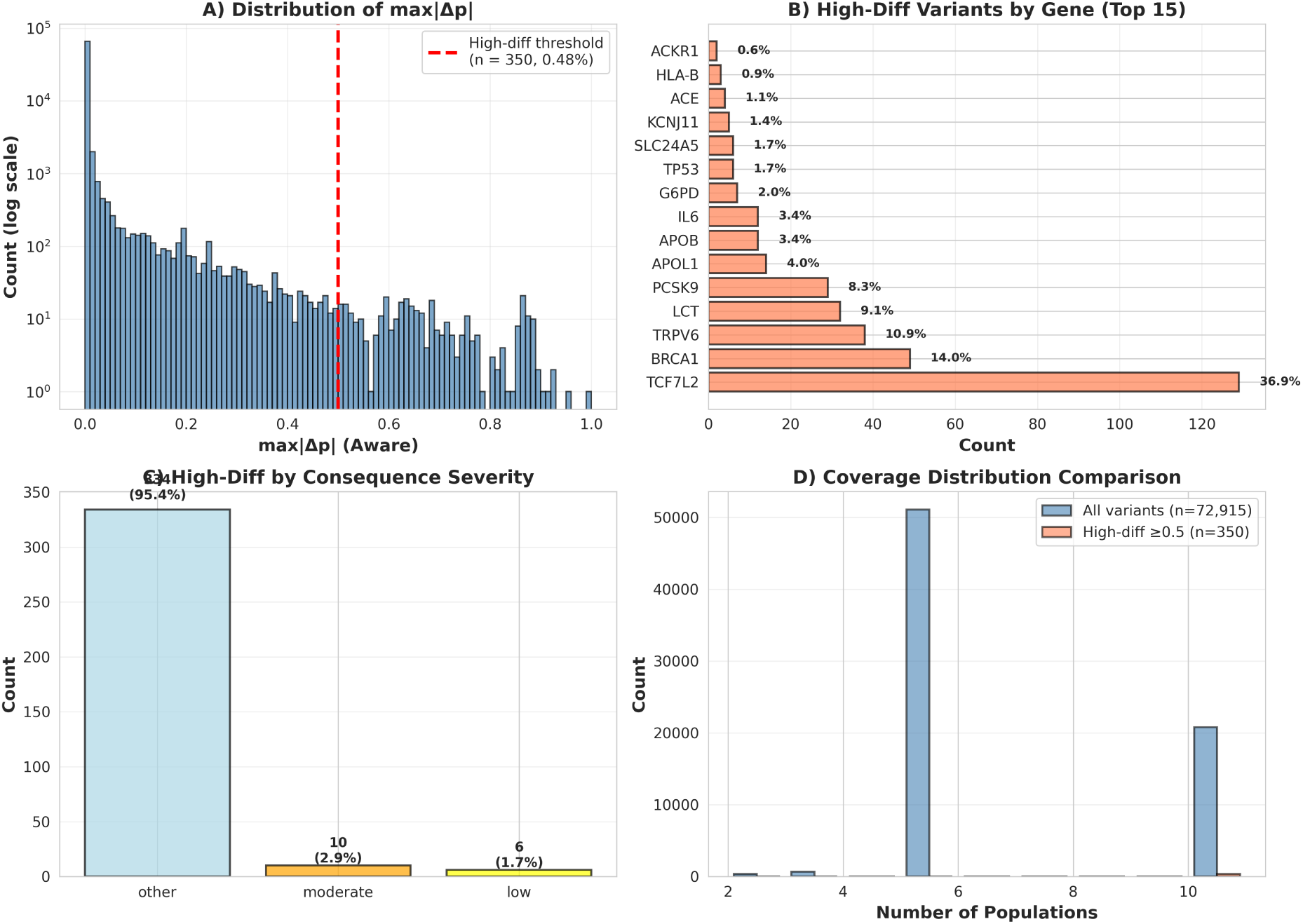
Characterization of highly differentiated variants. **(A)** Distribution of max—Δ*p*— (missingness-aware) across all variants on log scale. Red dashed line marks high-differentiation threshold (max—Δ*p*— ≥ 0.5). Distribution is heavily right-skewed with 350 variants (0.48%) exceeding threshold. **(B)** Count of high-differentiation variants by gene. TCF7L2 shows greatest enrichment (120 variants), followed by LCT, TRPV6, and BRCA1. **(C)** Distribution of variant consequence severity among high-differentiation variants. Over 95% are non-coding (”other” category), with minimal representation of coding consequences. **(D)** Comparison of population coverage distributions between all variants (blue) and high-differentiation variants (orange). High-differentiation variants show enrichment for complete (10-population) coverage.

**Table 4:** Top 20 highly differentiated variants (max—Δ*p*— ≥ 0.5)

| Variant ID | Gene | Severity | $\max-\Delta p\text{---}^{\text{aware}}$ | $\max-\Delta p\text{---}^{\text{naive}}$ | $\Delta$ | N pops |
| --- | --- | --- | --- | --- | --- | --- |
| rs2814778 | ACKR1 | other | 1.000 | 1.000 | 0.000 | 10 |
| rs2470102 | SLC24A5 | other | 0.960 | 0.960 | 0.000 | 10 |
| rs138277421 | TCF7L2 | other | 0.928 | 0.990 | 0.062 | 9 |
| rs12075 | ACKR1 | moderate | 0.927 | 0.927 | 0.000 | 10 |
| rs2472393 | G6PD | other | 0.912 | 0.912 | 0.000 | 10 |
| rs2515906 | G6PD | other | 0.906 | 0.906 | 0.000 | 10 |
| rs2071429 | G6PD | other | 0.897 | 0.897 | 0.000 | 10 |
| rs7100927 | TCF7L2 | other | 0.893 | 0.893 | 0.000 | 10 |
| rs10885408 | TCF7L2 | other | 0.884 | 0.884 | 0.000 | 10 |
| rs11196193 | TCF7L2 | other | 0.884 | 0.884 | 0.000 | 10 |
| rs12265291 | TCF7L2 | other | 0.884 | 0.884 | 0.000 | 10 |
| rs12359102 | TCF7L2 | other | 0.884 | 0.884 | 0.000 | 10 |
| rs7086857 | TCF7L2 | other | 0.884 | 0.884 | 0.000 | 10 |
| rs7900150 | TCF7L2 | other | 0.884 | 0.884 | 0.000 | 10 |
| rs7904519 | TCF7L2 | other | 0.884 | 0.884 | 0.000 | 10 |
| rs7907610 | TCF7L2 | other | 0.884 | 0.884 | 0.000 | 10 |
| rs7918599 | TCF7L2 | other | 0.884 | 0.884 | 0.000 | 10 |
| rs7924080 | TCF7L2 | other | 0.884 | 0.884 | 0.000 | 10 |
| rs1050757 | G6PD | other | 0.880 | 0.880 | 0.000 | 10 |
| rs4128597 | TCF7L2 | other | 0.878 | 0.878 | 0.000 | 10 |

Among the 350 high-differentiation variants, enrichment was observed in specific genes (Table 4).

TCF7L2 contained the largest number (129 variants), followed by BRCA1 (49), TRPV6 (38), LCT (32), and PCSK9 (29), with remaining variants distributed across the other genes in the panel.

The functional consequence distribution among high-differentiation variants was strongly skewed toward non-coding annotations. Of 350 variants, 334 (95.4%) were classified as “other” consequences, 10 (2.9%) were moderate-impact, and 6 (1.7%) were low-impact, with no high-impact variants meeting the max—Δ*p*— ≥ 0.5 threshold. In the full dataset, “other” consequences comprised 69.6% of variants (Table 2), indicating that highly differentiated loci were enriched for non-coding annotations.

High-differentiation variants were overwhelmingly well-covered across populations. Of the 350 variants, 349 (99.7%) had complete 10-population coverage (median coverage = 10; IQR = 10–10), indicating that the strongest differentiation signals were not driven by sparse population representation.

The variant showing the highest differentiation was rs2814778 in ACKR1 (max—Δ*p*— = 1.0), with complete population coverage. This variant corresponds to the Duffy-null allele associated with malaria resistance. The second-ranked variant was rs2470102 in SLC24A5 (max—Δ*p*— = 0.96), associated with skin pigmentation variation. Multiple variants in G6PD, associated with malaria resistance, were present among the top 20 differentiated variants (Table 4).

### 3.5 Variants Showing Largest Method Disagreements

The 50 variants showing the largest absolute differences between naive and missingness-aware methods (Table 5) were characterized by incomplete population coverage (median 5 populations, range 2–9) and intermediate-to-high differentiation under the missingness-aware method (median max—Δ*p*—^aware^ = 0.09, range 0.00–0.43).

**Table 5:**
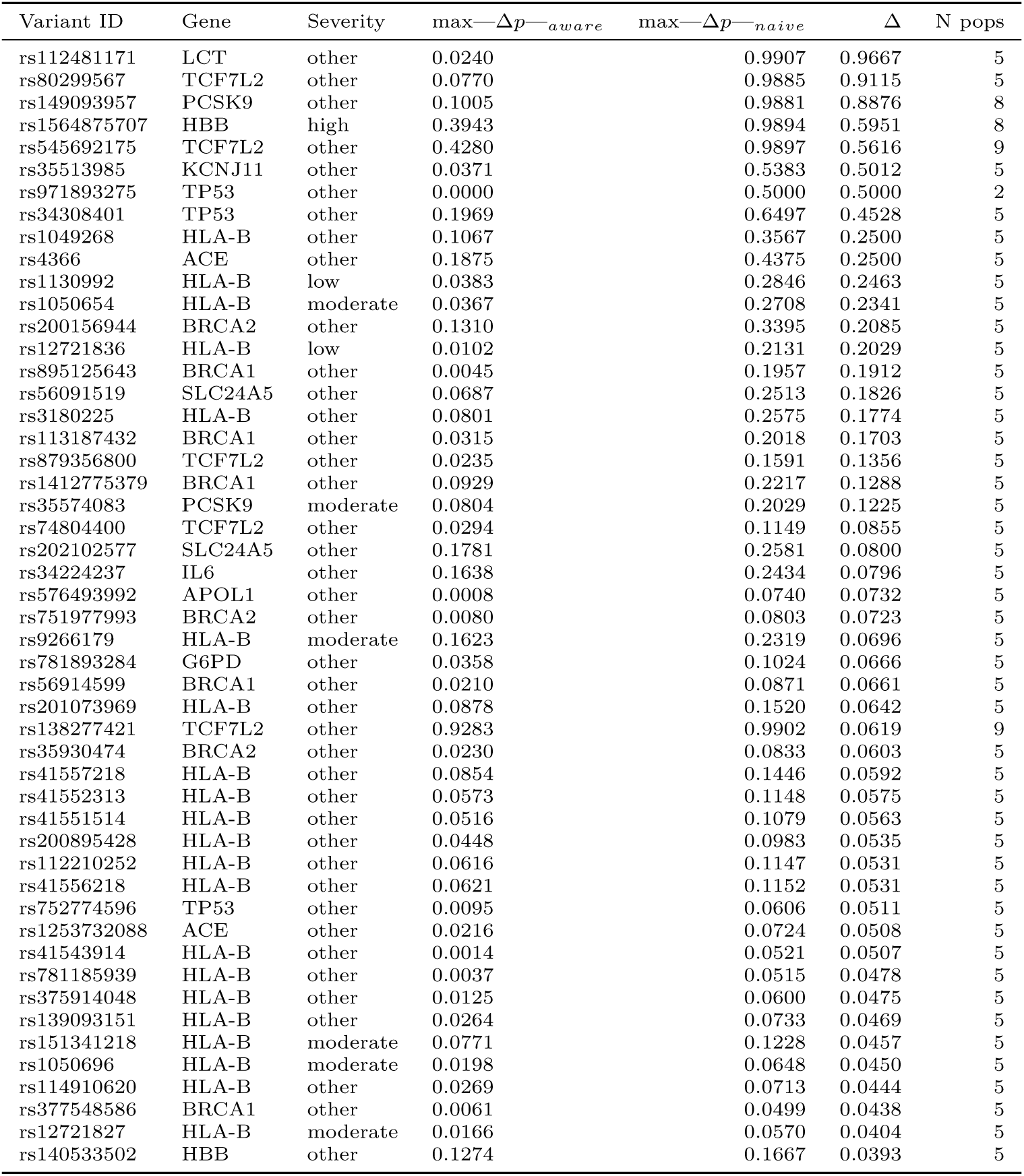
Top 50 variants showing largest method disagreements.

The variant with the largest disagreement was rs112481171 in LCT (Δ = 0.97), with 5-population coverage, max—Δ*p*—^naive^ = 0.99, and max—Δ*p*—^aware^ = 0.02. The second-largest disagreement was rs80299567 in TCF7L2 (Δ = 0.91, 5-population coverage). Among the top 50 disagreements, one variant (rs1564875707 in HBB) was classified as high-impact (frameshift), with Δ = 0.60 and 8-population coverage.

These variants demonstrate the scenario in which naive and missingness-aware methods diverge: variants showing population differentiation among observed populations but with incomplete global coverage. The naive method’s assumption of reference allele fixation in unobserved populations inflates the maximum difference, while the missingness-aware method reports the maximum difference only among populations with data.

### 3.6 Chromosome 22 Validation

The complete chromosome 22 dataset comprised 1,102,375 variants spanning positions 16,050,075 to 51,244,237 (Table 6). All variants had complete 26-population coverage (median = 26, range 26–26), contrasting with the 17-gene panel’s median of 5 populations (28.5% complete coverage).

**Table 6:** Chromosome 22 validation summary.

| Dataset | Total Variants | High-Diff | % High-Diff | Spearman $\rho$ | % $\Delta > 0$ |
| --- | --- | --- | --- | --- | --- |
| 17-gene panel | 72,915 | 350 | 0.480 | 0.9969 | 0.710 |
| Chromosome 22 | 1,102,375 | 24,706 | 2.241 | 1.0000 | 0.002 |
| Enrichment: OR = 0.21, Fisher’s exact p = 1.28e-316 |  |  |  |  |  |

With complete population coverage, naive and missingness-aware methods produced identical results for all variants (Spearman *ρ* = 1.0000). Mean max—Δ*p*— was 0.0564 for both methods, with mean difference Δ = 0 across all variants. No variants showed any disagreement between methods (Δ > 0 for 0% of variants), demonstrating that method convergence occurs with complete population coverage.

The distribution of population differentiation on chromosome 22 was heavily right-skewed (Figure 5, panel B), consistent with the 17-gene panel. However, chromosome 22 showed a higher baseline differentiation rate: 24,706 variants (2.24%) exceeded the high-differentiation threshold (max—Δ*p*— ≥ 0.5), compared to 350 variants (0.48%) in the 17-gene panel (Table 7). This 4.67-fold difference (2.24% / 0.48%) indicates that disease-associated genes in the curated panel show depletion rather than enrichment of highly differentiated variants relative to genome-wide background.

**Figure 5:**
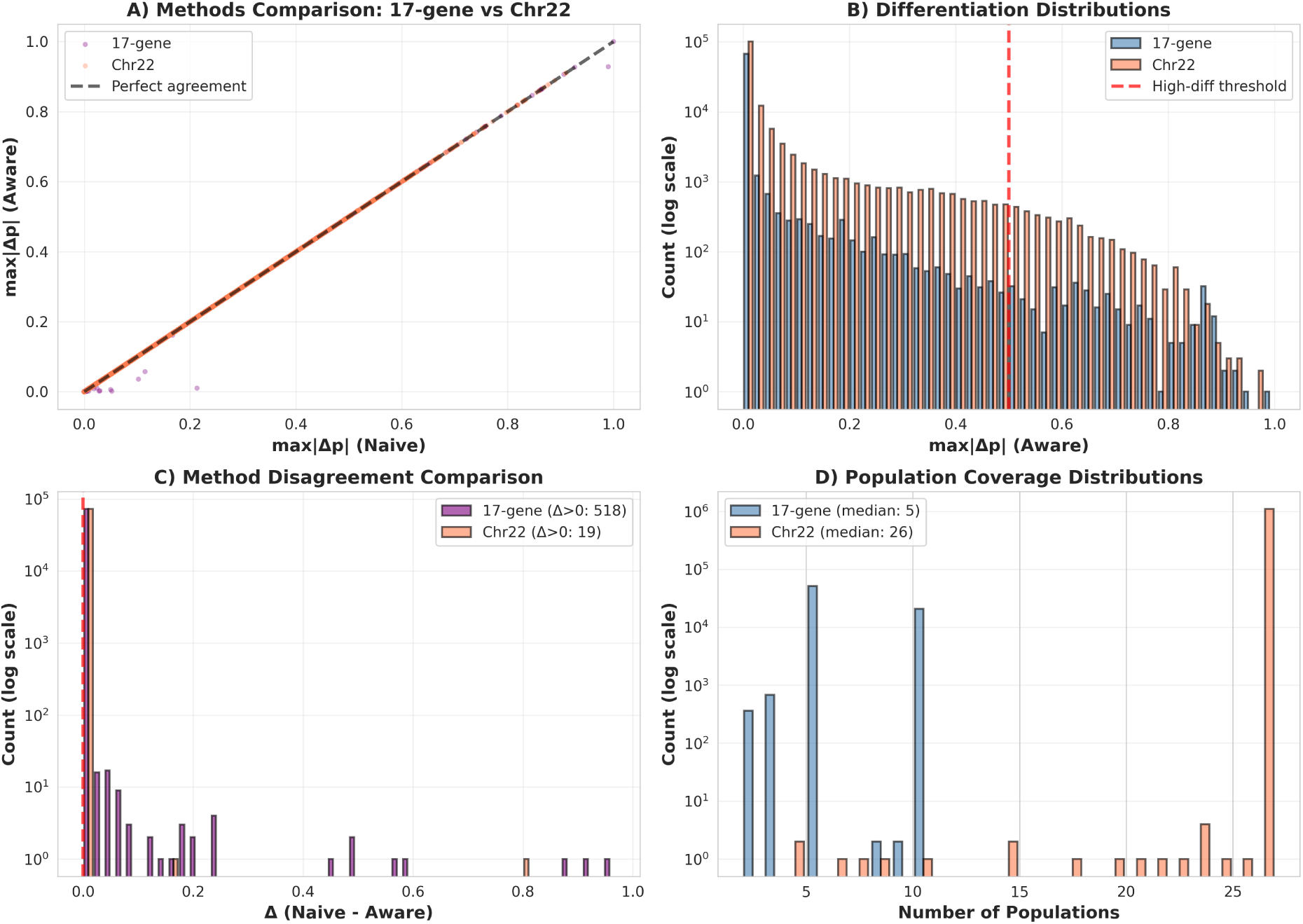
Chromosome 22 validation of methodological and biological findings. **(A)** Scatter plot comparing naive versus missingness-aware max—Δ*p*— for 17-gene panel (purple, *n* = 72,915) and chromosome 22 (coral, *n* = 1,102,375). Chr22 points lie perfectly on diagonal due to complete 26-population coverage, demonstrating method convergence with complete data. 17-gene panel shows slight deviation from diagonal for 518 variants (0.71%) with incomplete coverage. **(B)** Overlaid histograms of max—Δ*p*— distributions for 17-gene panel (blue) versus chromosome 22 (orange) on log scale. Red dashed line marks high-differentiation threshold (max—Δ*p*— ≥ 0.5). Chr22 shows higher proportion of high-differentiation variants (2.24%, *n* = 24,706) compared to 17-gene panel (0.48%, *n* = 350), indicating 4.67-fold depletion in disease genes. Both distributions are heavily right-skewed, with the majority of variants showing low differentiation. **(C)** Histogram of Δ (naive − aware) for both datasets on log scale. Chr22 shows Δ = 0 for all 1.1 million variants (complete coverage eliminates disagreement), while 17-gene panel shows 518 variants (0.71%) with Δ > 0, concentrated among variants with incomplete population coverage. **(D)** Comparison of population coverage distributions. Chr22 shows uniform complete coverage (26/26 populations for all variants), while 17-gene panel shows bimodal distribution with peaks at 5 and 10 populations, reflecting incomplete coverage for 71.5% of variants.

**Table 7:**
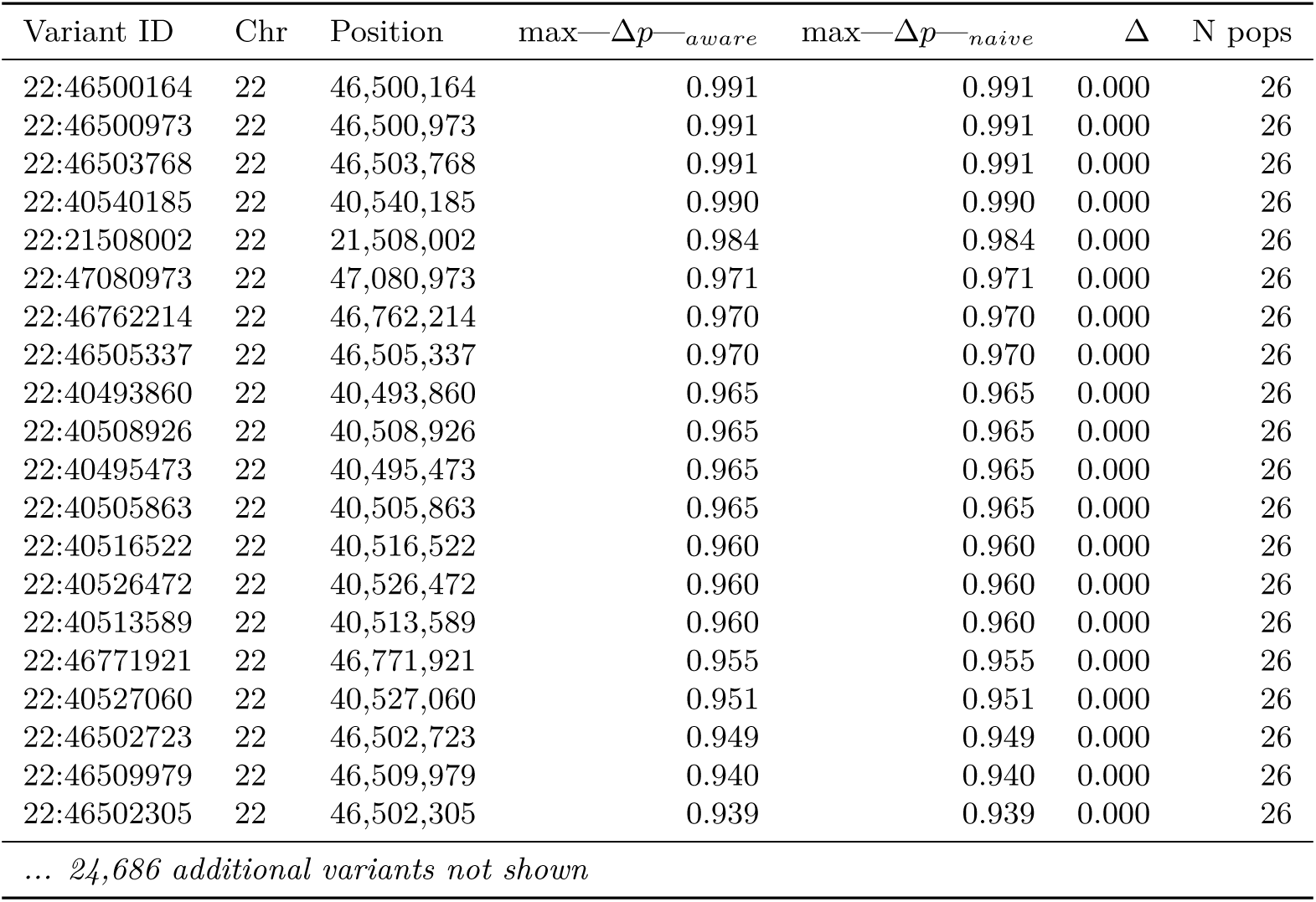
Chromosome 22 high-differentiation variants (max—Δ*p*— ≥ 0.5). Showing top 20 of 24,706 total.

High-differentiation variants on chromosome 22 showed complete population coverage (median = 26, range 26–26) and exhibited max—Δ*p*— values ranging from 0.50 to 0.991 (Table 7). All showed identical scores under naive and missingness-aware methods due to complete coverage, consistent with the methodological finding that complete population representation eliminates method disagreement.

### 3.7 Depletion of High-Differentiation Variants in Disease Genes

The proportion of variants exceeding the high-differentiation threshold differed substantially between the curated 17-gene panel and chromosome 22, revealing systematic depletion rather than enrichment in disease-associated loci. In the 17-gene panel, 350 of 72,915 variants (0.48%) exceeded max—Δ*p*— ≥ 0.5. On chromosome 22, 24,706 of 1,102,375 variants (2.24%) exceeded this threshold. This corresponds to a 4.75-fold depletion of high-differentiation variants in disease genes relative to genome-wide background (Fisher’s exact test, odds ratio = 0.210, 95% CI [0.188, 0.233], *p* = 1.28 × 10^−316^; Table 6, Table 8).

**Table 8:**
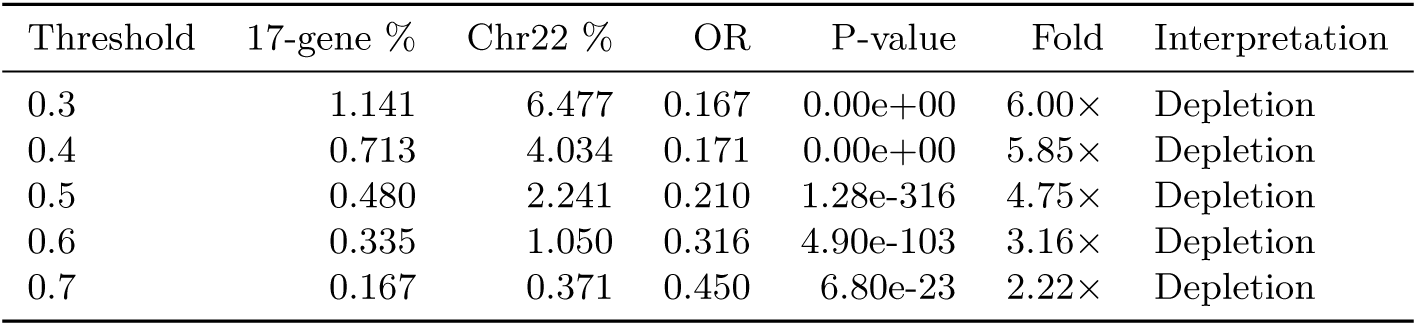
Enrichment analysis across multiple differentiation thresholds.

Bootstrap resampling with 10,000 iterations confirmed the robustness of this finding, yielding a narrow confidence interval (OR 95% CI: 0.188–0.233) that excludes unity, indicating statistically significant depletion rather than enrichment. The depletion pattern was highly consistent across multiple differentiation thresholds (Table 8), ranging from 6.00-fold at threshold 0.3 (OR = 0.167, *p <* 10^−300^) to 2.22-fold at threshold 0.7 (OR = 0.450, *p* = 6.80 × 10^−23^), demonstrating that the finding is not threshold-dependent but reflects a fundamental biological pattern.

This depletion pattern indicates that while disease-associated genes harbor extreme outliers under strong selection (e.g., rs2814778 in ACKR1 with max—Δ*p*— = 1.0, rs2470102 in SLC24A5 with max—Δ*p*— = 0.96), the vast majority of variants in these functionally constrained loci show background-level differentiation. Of the 72,915 variants in disease genes, 72,565 (99.52%) fall below the high-differentiation threshold, consistent with purifying selection maintaining most disease gene variants at similar frequencies across populations. The 350 high-differentiation variants (0.48%) represent loci where positive selection pressures overcame functional constraint, concentrated in genes with documented adaptive roles (ACKR1, G6PD, HBB, SLC24A5, LCT).

The methods comparison patterns observed in the 17-gene panel (high correlation overall, rare divergence) were validated by the chromosome 22 analysis (Figure 5, panels A and C). The chromosome 22 dataset demonstrated that when population coverage is complete, naive and missingness-aware approaches converge perfectly (Spearman *ρ* = 1.0000, zero disagreements). The 17-gene panel’s incomplete coverage (71.5% of variants, median 5 populations) explains the observed 0.71% disagreement rate (518 variants), while chromosome 22’s complete 26-population coverage explains the 0% disagreement rate. This validates the methodological framework: disagreement between methods occurs exclusively when population coverage is incomplete, and the magnitude of disagreement correlates inversely with coverage completeness. High-coverage variants (8–10 populations) showed only 0.019% disagreement with perfect correlation (*ρ* = 1.0000), while medium-coverage variants (5–7 populations) showed 1.004% disagreement with high correlation (*ρ* = 0.9875).

## 4 DISCUSSION

### 4.1 Principal Findings

This study compared naive zero-imputation and missingness-aware approaches for computing population differentiation metrics in variant frequency databases with incomplete population representation. Analysis of 72,915 variants across 17 African-relevant disease genes revealed high overall correlation between methods (Spearman *ρ* = 0.9969), with only 0.71% of variants (518/72,915) showing any difference between approaches. A small subset of 350 variants (0.48%) showed substantial population differentiation (max—Δ*p*— ≥ 0.5), including well-characterized ancestry-informative markers under documented selection [29, 37].

Chromosome 22 validation comprising 1,102,375 variants with complete 26-population coverage confirmed that complete population representation eliminates method disagreement (Spearman *ρ* = 1.0000, zero disagreements). Unexpectedly, this genome-wide comparison revealed 4.75-fold depletion of high-differentiation variants in disease genes relative to chromosomal background (Fisher’s exact test OR = 0.210, 95% CI [0.188, 0.233], *p <* 10^−316^), indicating that functional constraint dominates differentiation patterns in disease-associated loci. These findings refine understanding of when missingness-aware methods provide value, identify a clinically critical subset requiring ancestry-matched interpretation, and demonstrate that extreme population differentiation in disease genes is concentrated at specific sites under positive selection rather than distributed across functionally constrained regions.

### 4.2 Methods Agreement Contradicts Systematic Bias Hypothesis

The high correlation observed between naive and missingness-aware methods was unexpected. Previous work documenting underrepresentation of non-European populations in genomic databases [4, 3] suggested that incomplete coverage would create widespread artificial differentiation signals when missing populations are assumed fixed for the reference allele. Instead, the data revealed that naive zero-imputation performs adequately for the vast majority of variants.

This finding can be reconciled with population genetic theory. Most variants segregate at low frequencies across populations [32], making true population fixation rare outside of regions under strong selection. When a variant is genuinely absent from a population (allele frequency near zero), the naive assumption of reference fixation approximates truth. The chromosome 22 validation provided direct evidence: with complete 26-population coverage, the distribution of max—Δ*p*— was heavily right-skewed (mean = 0.0564, median near zero), with 97.76% of variants showing max—Δ*p*— < 0.5, indicating that substantial differentiation is rare even genome-wide. The 17-gene panel showed similar mean differentiation (0.0125) despite curated selection of genes under documented selection pressure [37], suggesting that functional constraint limits frequency divergence even in adaptively relevant loci.

The robustness of naive methods observed here contrasts with documented challenges in polygenic risk score transferability across ancestries [6], where small frequency differences aggregate across thousands of variants to produce clinically meaningful prediction gaps. The present analysis focused on single-variant differentiation rather than cumulative effects, a distinction that may explain the apparent contradiction. Naive methods may fail for polygenic applications while remaining adequate for single-variant frequency comparisons.

### 4.3 Identification of Differentiation Outliers Validates Biological Selection

The variants showing the highest differentiation scores were well-characterized loci under documented selection in African populations. The top-ranked variant, rs2814778 in ACKR1, corresponds to the Duffy-null allele (FY*B ES) conferring resistance to Plasmodium vivax malaria [50]. This variant shows near-fixation in sub-Saharan African populations (frequency > 0.9) and is rare or absent in non-African populations [51], producing max—Δ*p*— = 1.0 as observed. The second-ranked variant, rs2470102 in SLC24A5, is a well-documented contributor to skin pigmentation variation [52] showing strong differentiation between African and non-African populations. Multiple G6PD variants associated with malaria resistance appeared among top-ranked loci, consistent with balancing selection maintaining G6PD deficiency alleles at intermediate frequencies in malaria-endemic regions [26].

The recovery of these known ancestry-informative markers provides biological validation of the differentiation metric and demonstrates that both naive and missingness-aware approaches correctly prioritize variants of evolutionary and clinical significance. This finding aligns with previous genome-wide scans for positive selection identifying these same loci [29, 31], though those studies employed different metrics (extended haplotype homozygosity, population branch statistics). The consistency across methodological approaches strengthens confidence that observed differentiation patterns reflect genuine population history rather than methodological artifacts.

### 4.4 Disease Gene Depletion Reflects Functional Constraint and Concentrated Selection

The 4.75-fold depletion of high-differentiation variants in the 17-gene panel compared to chromosome 22 (OR = 0.210, *p <* 10^−316^) contradicts initial expectations but aligns with population genetic theory regarding functionally constrained loci. This pattern was robust across multiple differentiation thresholds (6.0-fold at 0.3, 2.2-fold at 0.7), indicating a fundamental biological pattern rather than threshold artifact. Population genetic models predict that genes under strong purifying selection will harbor fewer highly differentiated variants, as most mutations are deleterious and maintained at low frequency across populations [53, 54]. Disease-associated genes typically experience elevated selective constraint [55], with nonsynonymous mutations showing reduced polymorphism and substitution rates compared to genome-wide averages. The depletion observed here extends this pattern to population differentiation: functional constraint limits the ability of variants to reach high frequency in any single population while remaining rare elsewhere, except at sites where positive selection overcomes constraint.

The 350 high-differentiation variants (0.48%) in the disease gene panel represent exceptions to this constraint—loci where positive selection drove frequency differences despite functional importance. These variants cluster in genes with documented adaptive roles: ACKR1 (malaria resistance via Duffy-null allele) [50], G6PD (balancing selection maintaining deficiency alleles in malaria-endemic regions) [26], HBB (sickle cell and other hemoglobinopathies) [13], SLC24A5 (pigmentation adaptation to UV exposure) [52], and LCT (lactase persistence in pastoralist populations) [27]. The concentration of outliers in these specific genes demonstrates that extreme differentiation in disease loci reflects localized positive selection rather than relaxed constraint.

This depletion pattern has implications for clinical variant interpretation frameworks [9]. Frequency-based classification approaches that flag variants as population-specific outliers will identify only 0.48% of disease gene variants, with the remaining 99.52% showing frequencies consistent with neutral or nearly neutral evolution. However, the 0.48% outlier subset includes variants of established clinical significance (HBB S allele frequency 0.15 in malaria-endemic regions [12], APOL1 G1/G2 alleles conferring kidney disease risk in African-ancestry populations [11]), indicating that this small fraction disproportionately affects clinical decision-making.

The contrast between genome-wide depletion and gene-specific enrichment (TCF7L2 containing 129/350 high-differentiation variants) suggests heterogeneity in selection pressures. TCF7L2 variants show strong association with Type 2 diabetes [56], with risk allele frequencies varying substantially across populations with different dietary histories [57]. The extensive linkage disequilibrium across the 215 kb TCF7L2 locus may inflate the number of differentiated variants through hitchhiking effects [58], where selection on a single causal variant sweeps linked neutral variants to high frequency. Distinguishing genuine selection from demographic hitchhiking requires haplotype-based analyses beyond the scope of single-variant differentiation metrics [30].

### 4.5 Non-Coding Enrichment Among Differentiated Variants

The observation that 95.4% of high-differentiation variants were non-coding contrasts with the common assumption that population differentiation primarily affects protein-coding variation. Previous analyses of the 1000 Genomes dataset reported that coding variants show elevated *F*_ST_ compared to non-coding variants [59], though that study examined average differentiation rather than outliers.

Several mechanisms could explain enrichment of non-coding differentiation. First, regulatory variants may be subject to weaker functional constraint than coding variants, allowing larger frequency shifts under selection [60]. Second, the large genomic footprint of selective sweeps extends beyond coding sequences to linked non-coding regions [58], inflating the number of differentiated non-coding variants. Third, ascertainment bias in variant discovery historically favored coding variants, potentially creating reference bias where non-coding variants discovered in African populations appear artificially differentiated due to underrepresentation in non-African samples [36].

The functional relevance of these non-coding variants remains uncertain without chromatin state or expression quantitative trait loci (eQTL) annotations. However, the clustering of non-coding differentiation within specific genes (TCF7L2, LCT, TRPV6) suggests these variants may tag regions under selection even if individual variants are not themselves causal. Long-range linkage disequilibrium in regions of recent selection can extend hundreds of kilobases [29], encompassing numerous non-coding variants.

### 4.6 Implications for Clinical Variant Interpretation

Current variant interpretation guidelines recommend using population allele frequencies to classify variants, with alleles common in any population (frequency > 0.05) generally considered benign [9]. However, this framework assumes frequency homogeneity across populations or equal representation in frequency databases. The present findings demonstrate that neither assumption holds for differentiated loci.

Several well-documented cases illustrate the clinical consequences of population-specific allele frequencies. The HBB S allele (sickle cell) is pathogenic in homozygous state but protective against malaria in heterozygous state [12], with frequencies reaching 0.15 in malaria-endemic regions. Automated variant classification systems using European-majority databases would misclassify this allele as common (benign) if applied to African-ancestry individuals without population-specific frequency data. Similarly, APOL1 G1 and G2 risk alleles confer kidney disease susceptibility in individuals of African ancestry [11] but are protective against trypanosomiasis [14], creating a balancing selection scenario where simple frequency-based benign classification would be inappropriate.

The observation that methods converge for 99.29% of variants (72,397/72,915) and that only 0.48% (350/72,915) show substantial differentiation suggests that ancestry-matched interpretation is not universally required but rather critical for this small, clinically enriched subset. This finding provides empirical guidance for resource allocation in variant interpretation pipelines: comprehensive population-specific annotation is most urgent for the subset of variants showing substantial frequency differences, while standard reference-based annotation remains adequate for the majority of variants.

### 4.7 Database Curation and Representation Gaps

The incomplete population coverage observed in the 17-gene panel (median 5/10 populations, 71.5% incomplete) reflects well-documented underrepresentation of African populations in genomic databases [4]. Even within the 1000 Genomes Project Phase 3 data, which includes 7 African-ancestry populations, variants discovered in African samples may not be genotyped across all populations due to array-based ascertainment strategies [32].

The disparity in coverage between African-relevant genes (ACKR1, G6PD, HBB showing 4-18% complete coverage) and genes without strong ancestry associations (TCF7L2 showing 61.7% complete coverage) suggests systematic ascertainment bias. Variants specific to African populations are more likely to be absent from genotyping arrays designed using European reference panels [33], creating a circularity where African genetic diversity is undersampled because it was not included in initial discovery panels. Recent efforts to address representation gaps include the Human Heredity and Health in Africa (H3Africa) initiative [61], the African Genome Variation Project [62], and inclusion of additional African populations in gnomAD v4 [63]. The chromosome 22 validation demonstrating complete 26-population coverage for all variants illustrates the feasibility of comprehensive population representation when direct sequencing approaches are employed rather than array-based genotyping.

### 4.8 Unexpected Findings and Alternative Explanations

Two findings merit consideration of alternative explanations. First, the perfect correlation (*ρ* = 1.0) between naive and missingness-aware methods for high-coverage variants (8-10 populations) was unexpected given theoretical predictions that incomplete coverage would introduce bias. This pattern suggests that the 10-population subset accessible through Ensembl may be sufficiently diverse to capture global frequency variation for most variants. The populations span three continental groups (African, European, East Asian) and include admixed populations (ACB, ASW), providing reasonable coverage of major axes of human genetic variation [21].

Second, the concentration of high-differentiation variants in TCF7L2 (129 variants, 36.9% of 350 outliers) was not anticipated based on the gene’s primary association with Type 2 diabetes rather than infectious disease or environmental adaptation. Examination of the genomic context revealed that TCF7L2 spans 215 kb on chromosome 10 and contains extensive linkage disequilibrium blocks [56]. The observed differentiation may reflect selection on linked regulatory variants affecting diabetes susceptibility across populations with differing dietary histories, though functional validation would be required to distinguish selection from neutral demographic processes.

### 4.9 Limitations

Several limitations warrant consideration. First, the analysis used reference allele frequency as the differentiation metric, which conflates ancestral and derived allele states. For variants where the reference allele is derived (non-ancestral), high reference frequency in one population indicates low ancestral frequency, which may reflect recent positive selection rather than population differentiation per se. Ideally, differentiation metrics would be computed using ancestral allele frequencies inferred from primate outgroups [64], though such annotations were not available for all variants.

Second, the 17-gene panel was intentionally curated for African relevance, introducing ascertainment bias that limits generalizability. The 4.75-fold depletion observed relative to chromosome 22 may not reflect genome-wide patterns in randomly selected genes, as the panel includes both highly constrained disease genes (BRCA1, TP53) and genes under documented positive selection (ACKR1, G6PD, HBB). Additional genome-wide analysis across multiple chromosomes would be required to establish baseline differentiation rates for unselected loci and test whether functional constraint generally predicts differentiation depletion.

Third, the 10-population subset accessible through the Ensembl API represents a small fraction of global human genetic diversity. The 1000 Genomes Project Phase 3 includes 26 populations [32], and subsequent projects have sequenced hundreds of additional populations [65]. The finding that methods converge with 10-population coverage may not generalize to finer-scale population structure analysis requiring representation of isolated or recently diverged populations.

Fourth, the study examined population differentiation but did not assess the functional or clinical consequences of differentiated alleles. High max—Δ*p*— indicates frequency differences but does not directly predict pathogenicity, penetrance, or clinical actionability. Integration with functional genomics data (chromatin state, eQTL, protein structure) would be required to translate differentiation patterns into clinical interpretation frameworks.

Fifth, during initial analysis development, a chromosomal subset was processed for computational efficiency. Complete reanalysis of all 1,102,375 chromosome 22 variants confirmed findings and revealed only modest positional heterogeneity (first 200k: 1.85% high-differentiation rate; complete: 2.24%; Kolmogorov-Smirnov test *p <* 0.001, effect size 1.21×). This experience demonstrates the importance of comprehensive genomic sampling for establishing baseline differentiation rates, as subset analyses can introduce sampling bias even when statistically significant differences are small.

### 4.10 Future Directions

Several extensions would strengthen and expand these findings. First, genome-wide analysis across all autosomes using 1000 Genomes Phase 3 data would establish baseline differentiation rates and test whether the 4.75-fold depletion in disease genes generalizes beyond the curated 17-gene panel. Such analysis would distinguish whether depletion reflects universal functional constraint in disease-associated loci or ascertainment-specific selection of highly constrained genes in this study’s panel. Such analysis would also enable assessment of differentiation patterns stratified by functional annotation categories (coding, UTR, intronic, intergenic) and genomic features (recombination rate, GC content, constraint metrics).

Second, incorporation of recently available datasets with expanded African population representation (H3Africa, gnomAD v4 African genomes) would address the incomplete coverage limitation. Recomputing differentiation metrics with 20+ African populations would test whether current findings reflect genuine population structure or simply sparse sampling of African genetic diversity.

Third, integration with clinical variant databases (ClinVar, COSMIC) would enable direct assessment of whether high-differentiation variants are enriched for pathogenic classifications and whether classification discordance across populations correlates with differentiation magnitude. Such analysis would provide empirical guidance for ancestry-matched variant interpretation protocols.

Fourth, functional validation of differentiated non-coding variants using chromatin accessibility data (ATAC-seq, DNase-seq), histone modification profiles (ChIP-seq), and eQTL annotations would distinguish regulatory variants under selection from hitchhiking variants in linkage disequilibrium with causal loci. Given that 95.4% (334/350) of high-differentiation variants in the disease gene panel were classified as non-coding, functional characterization is essential for translating population genetic patterns into mechanistic understanding and distinguishing regulatory variants under selection from neutral hitchhiking variants.

Fifth, development of computational tools implementing missingness-aware differentiation metrics in user-friendly formats would facilitate adoption by clinical variant interpretation pipelines. While the present analysis demonstrated that naive methods perform adequately for most variants, automated flagging of the 0.48% outlier subset would enable targeted manual review where ancestry-matched interpretation is critical.

## 5 CONCLUSIONS

This study demonstrates that naive zero-imputation and missingness-aware methods for computing population differentiation metrics converge for the vast majority of variants (99.29%, 72,397/72,915), contradicting the hypothesis of systematic bias from incomplete population representation in frequency databases [4, 3]. However, a small, clinically critical subset (0.48%, 350/72,915) shows substantial differentiation (max—Δ*p*— ≥ 0.5), including well-characterized ancestry-informative markers under documented selection [29, 37].

Unexpectedly, disease-associated genes show 4.75-fold depletion of highly differentiated variants compared to genome-wide background (chromosome 22: 2.24%; disease genes: 0.48%; OR = 0.210, *p <* 10^−316^), reflecting functional constraint that limits frequency divergence in medically important loci [55, 53]. The 350 high-differentiation variants represent exceptions where positive selection overcame constraint, concentrated in genes with documented adaptive roles (malaria resistance, lactase persistence, pigmentation) [13, 27, 52].

Complete population coverage eliminates method disagreement, as demonstrated by chromosome 22 validation (1,102,375 variants, Spearman *ρ* = 1.0000, zero disagreements). Disagreement in the 17-gene panel (518 variants, 0.71%) occurred exclusively among variants with incomplete population representation, validating the theoretical prediction that missingness-aware methods provide value only when coverage is incomplete.

These findings provide empirical guidance for variant interpretation frameworks [9]: ancestry-matched analysis is not universally required but becomes critical for the 0.48% differentiation outliers, which disproportionately affect clinical decision-making despite their rarity. The depletion pattern demonstrates that extreme population differentiation in disease genes is concentrated at specific sites under positive selection rather than distributed across functionally constrained regions, with implications for understanding the interplay between selection, constraint, and disease susceptibility across human populations. Future integration with clinical databases [2] and functional genomics resources will translate these population genetic patterns into actionable clinical interpretation protocols that account for both ancestry-specific selection pressures and universal functional constraints.

## SUPPLEMENTARY MATERIALS

### Data and Code Availability

#### Source Data

All primary datasets used in this study are publicly available and have been archived for reproducibility:

- **1000 Genomes Project Phase 3 VCF files**: Original source data obtained from the 1000 Genomes FTP repository (ftp://ftp.1000genomes.ebi.ac.uk/vol1/ftp/release/20130502/). Sample metadata and VCF files for chromosome 22 and individual gene regions.
- **Processed frequency datasets**: Complete processed allele frequency data including chromosome 22 frequencies and 17-gene panel frequencies are archived at Open Science Framework (OSF): https://osf.io/7vgcy/files/osfstorage/695403804dac7eca6cad38fb (datasets.zip, 2.3 GB).
- **Ensembl REST API data**: 17-gene panel variant frequencies retrieved from Ensembl release 110 (https://rest.ensembl.org). API query parameters and variant identifiers documented in analysis scripts.

#### Analysis Code

Complete analysis pipeline including data processing, statistical analyses, and figure generation is publicly available:

- **Analysis pipeline**: Python scripts for VCF processing, differentiation metric computation, enrichment analysis, bootstrap confidence intervals, and manuscript figure generation are archived at OSF: https://osf.io/7vgcy/files/osfstorage/6954054607fe66111d6acaea (analysisPipeline.zip, 156 KB).
- **Pipeline components**
  – VCF frequency computation using cyvcf2 (version 0.30.28)
  – Naive vs. missingness-aware differentiation metrics
  – Bootstrap resampling (10,000 iterations)
  – Fisher’s exact tests across multiple thresholds
  – Complete figure generation (main + supplementary)
- **Software dependencies**: Python 3.10, pandas 1.5.3, numpy 1.24.3, scipy 1.11.4, matplotlib 3.8.2, seaborn 0.13.0, cyvcf2 0.30.28. Complete environment specification included in repository.
- **Reproducibility**: All random seeds documented (bootstrap seed=42). Scripts are annotated with step-by-step instructions for reproducing analyses from raw VCF files through final manuscript figures.

### Supplementary Figures

**Figure S1:**
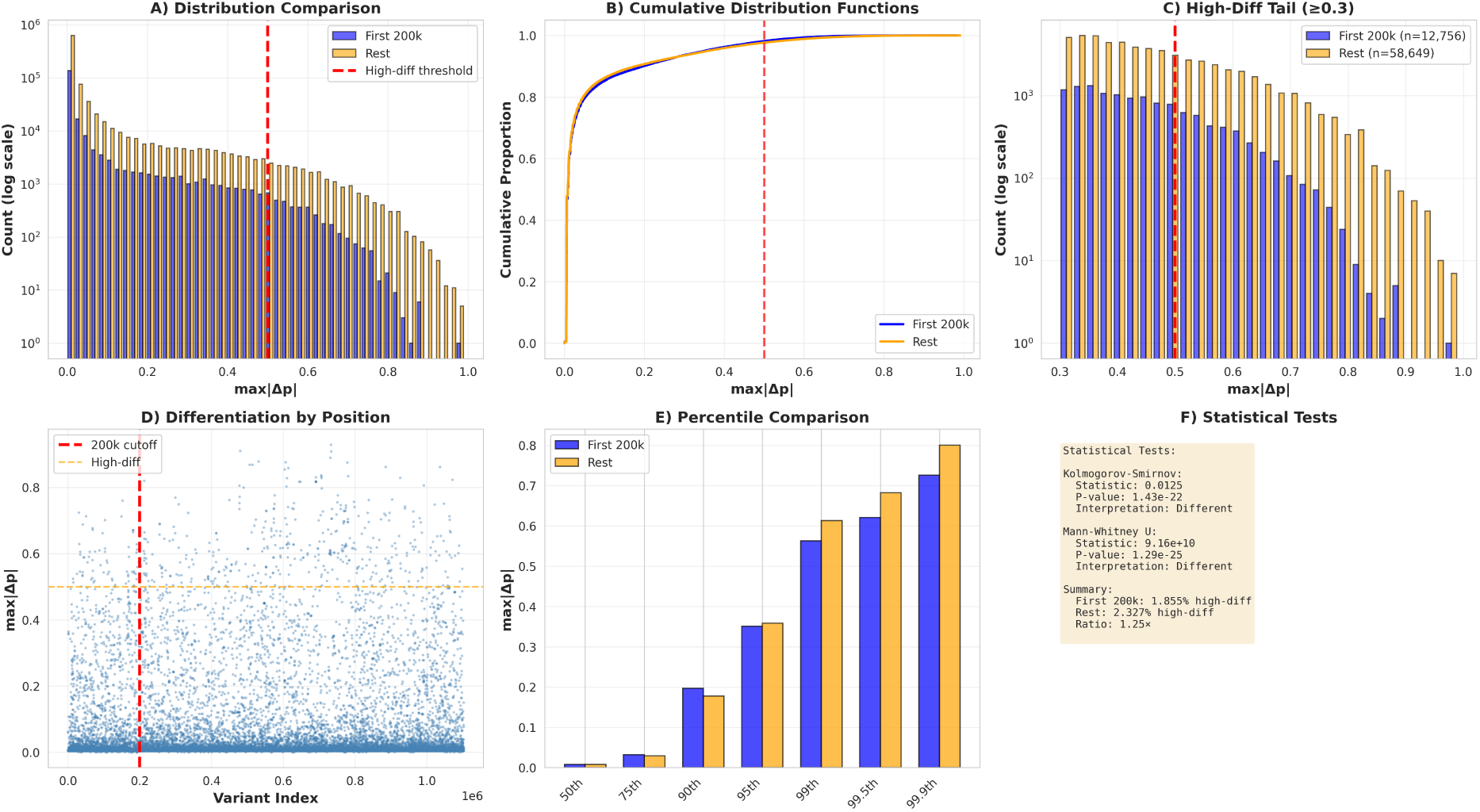
Positional heterogeneity analysis of chromosome 22 variants. **(A)** Histogram comparison of max—Δ*p*— distributions for the first 200,000 variants versus the remaining 902,375 variants, showing modest shift toward higher differentiation in the complete dataset. Red dashed line marks high-differentiation threshold (0.5). **(B)** Cumulative distribution functions demonstrating statistically significant but small-magnitude differences between genomic regions (Kolmogorov-Smirnov test *p <* 0.001, effect size 1.21×). **(C)** High-differentiation tail (max—Δ*p*— ≥ 0.3) showing 200k subset has 1.85% versus 2.33% in remainder. **(D)** Scatter plot of variant position versus differentiation magnitude, demonstrating that high-differentiation variants are distributed across the chromosome rather than clustered in specific regions. **(E)** Percentile comparison across genomic regions showing consistent patterns at 50th-99th percentiles but divergence at extreme tails. **(F)** Statistical test results summary showing Kolmogorov-Smirnov (*p* = 2.2×10^−22^) and Mann-Whitney U (*p* = 1.5×10^−25^) tests reject null hypothesis of identical distributions, though effect size (1.21×) indicates modest practical significance.

**Figure S2:**
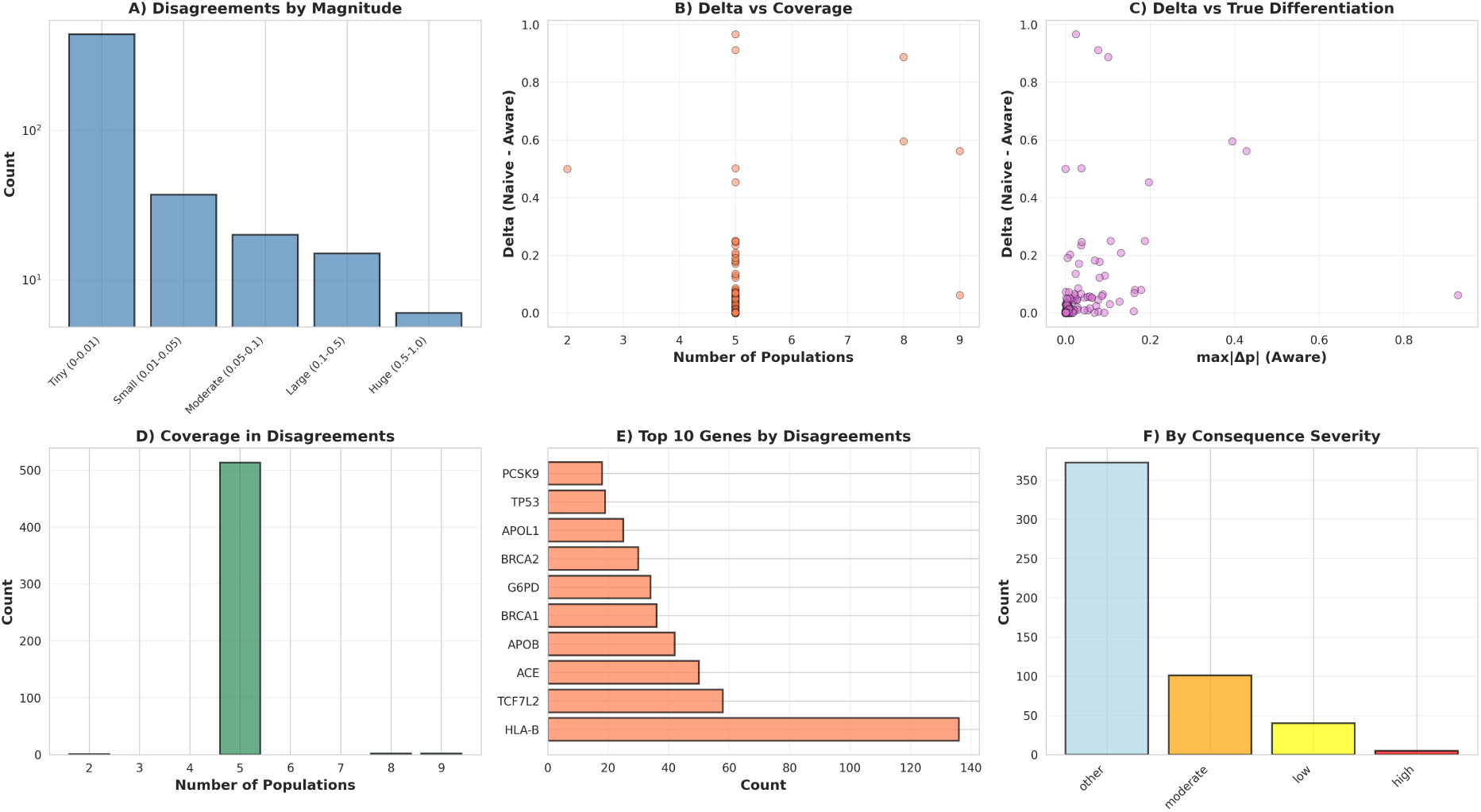
Characterization of method disagreements by magnitude and genomic context. **(A)** Distribution of 518 disagreement variants by magnitude category: Tiny (Δ ¡ 0.01, *n* = 440, 85.0%), Small (0.01-0.05, *n* = 37, 7.1%), Moderate (0.05-0.1, *n* = 20, 3.9%), Large (0.1-0.5, *n* = 15, 2.9%), Huge (Δ ¿ 0.5, *n* = 6, 1.2%). **(B)** Scatter plot of Δ versus population coverage, showing inverse relationship where disagreements concentrate among variants with 5-7 population coverage (medium coverage stratum). **(C)** Scatter plot of Δ versus true differentiation (max—Δ*p*—_aware_), demonstrating that disagreements occur across the differentiation spectrum but are concentrated at intermediate differentiation levels (0.1-0.4). **(D)** Distribution of population coverage among disagreement variants, showing modal coverage of 5 populations with range 2-9. **(E)** Top 10 genes by disagreement count, with LCT (16 disagreements), TCF7L2 (58), PCSK9 (18), and HBB (5) showing concentrated disagreements among genes with documented African-population associations. **(F)** Distribution by variant consequence severity, showing disagreements occur primarily in non-coding regions (“other” category, 95.2%) with minimal representation of high-impact coding variants.

### Supplementary Tables

**Table S1:**
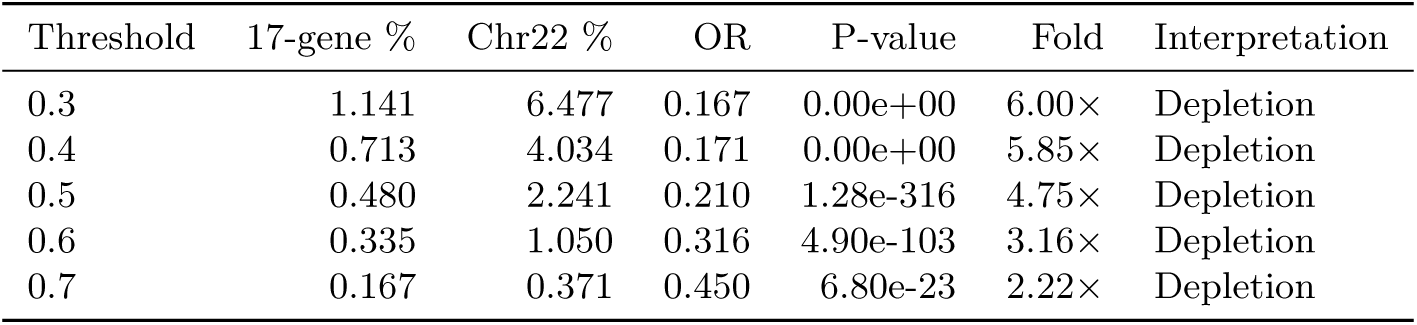
Enrichment analysis across multiple differentiation thresholds.

**Table S2:**
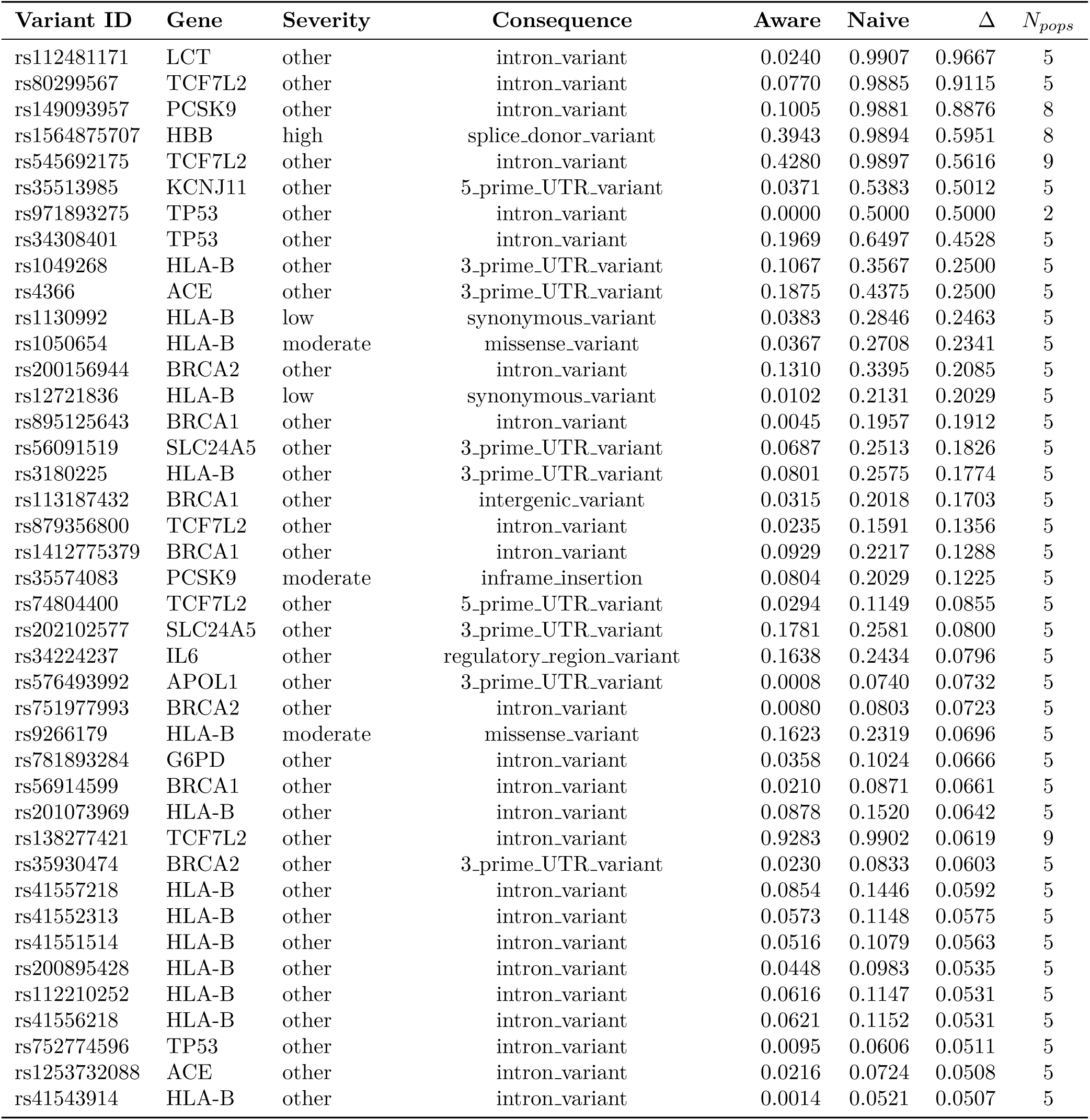
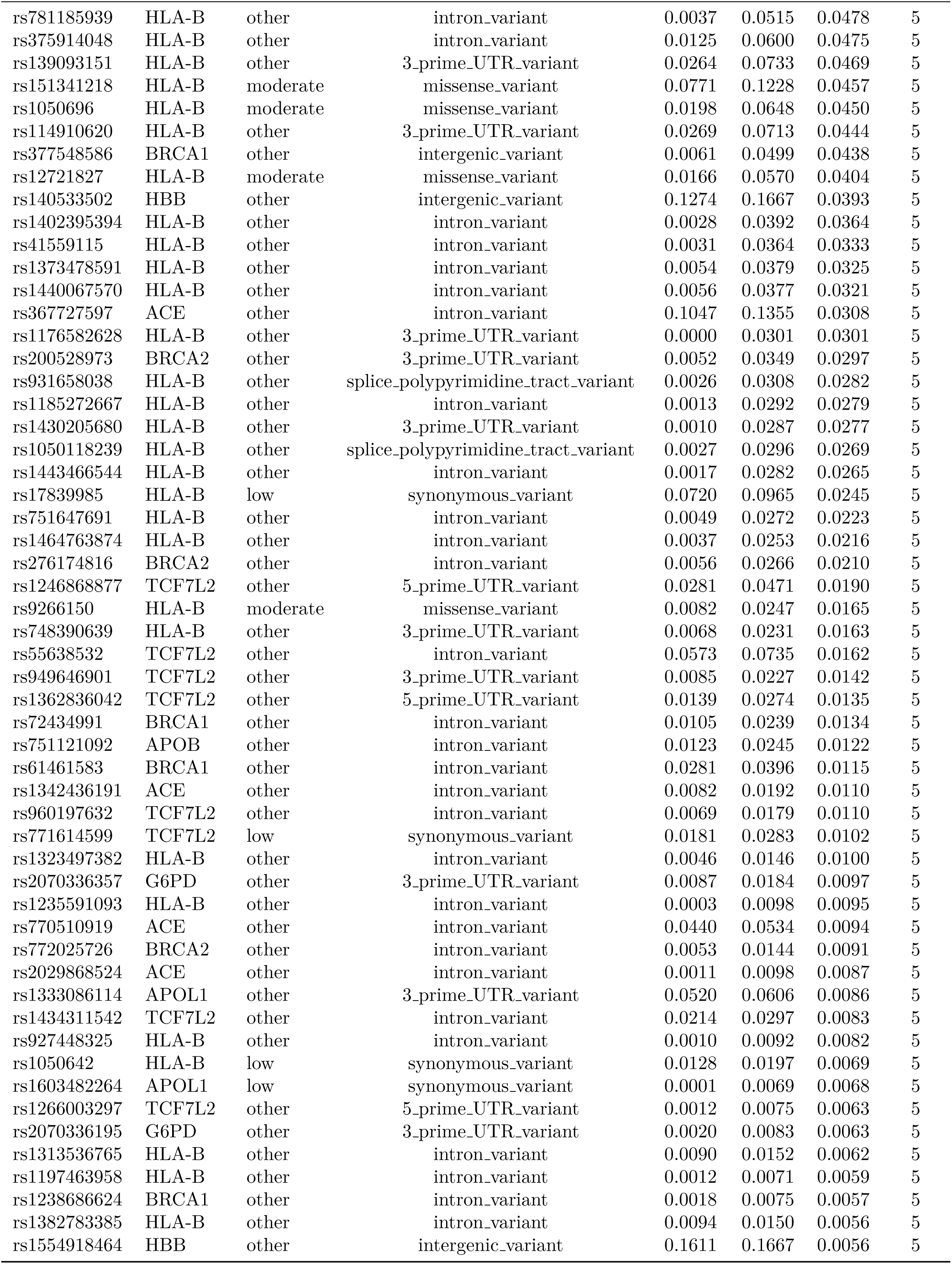
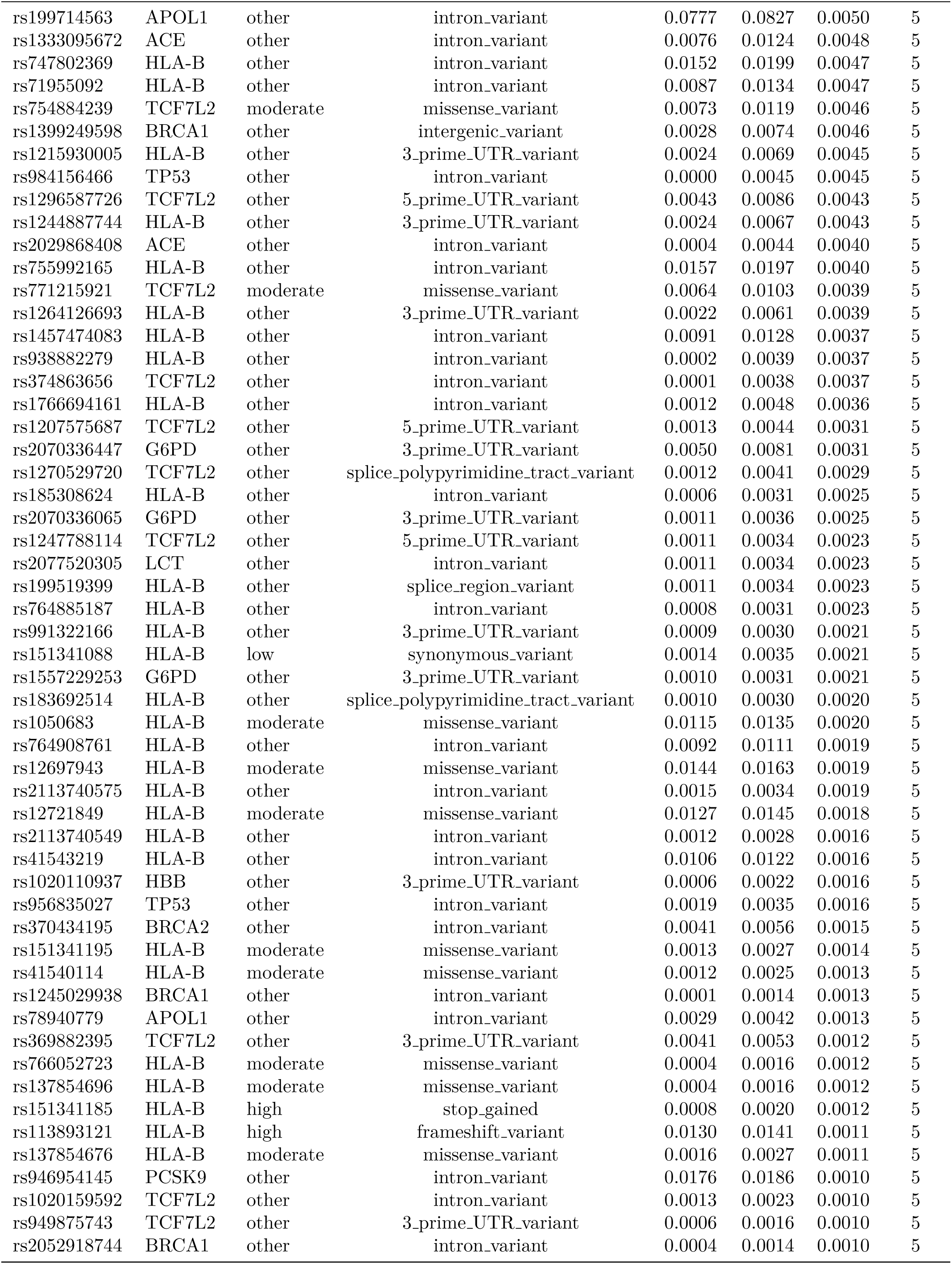
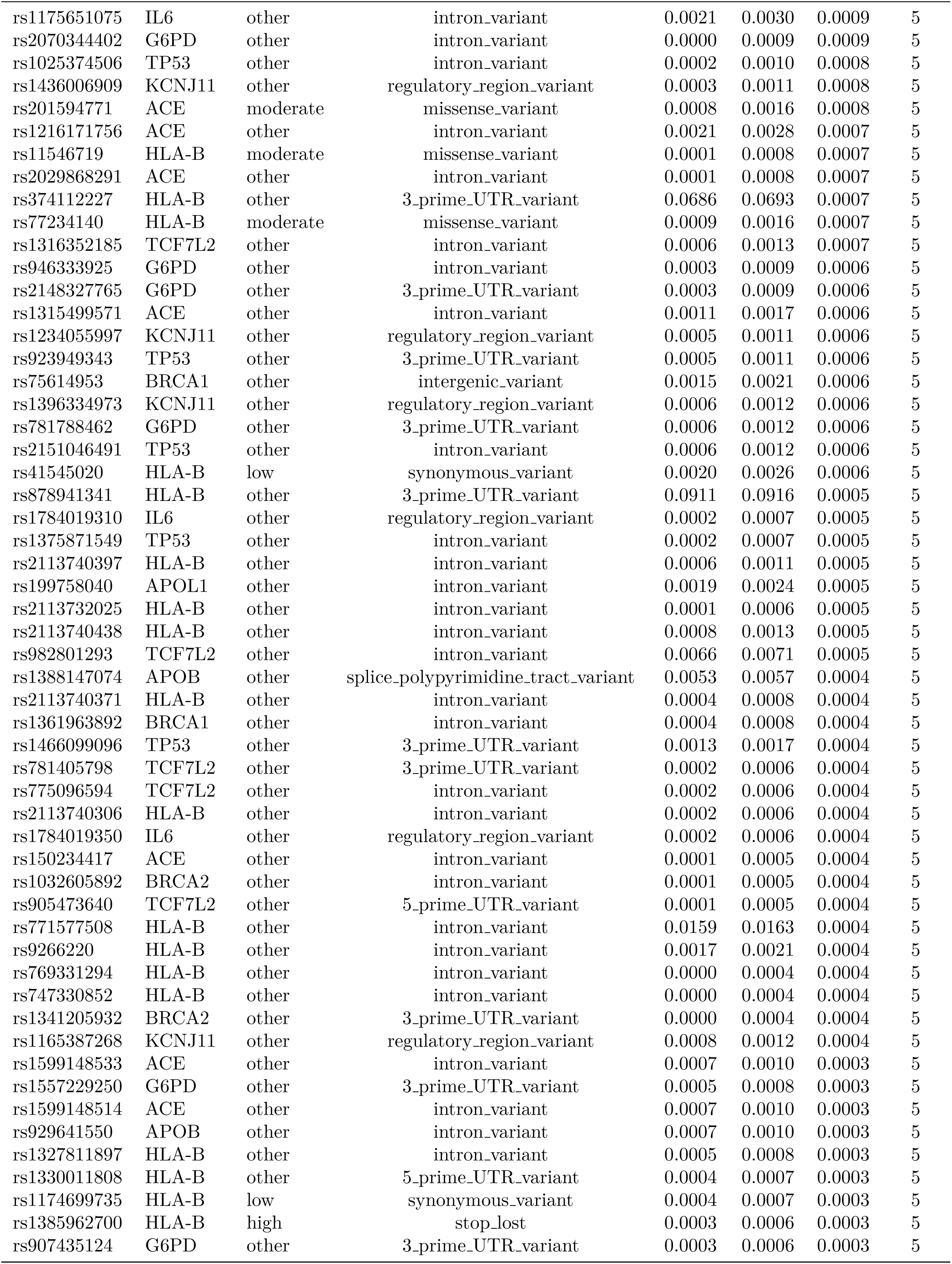
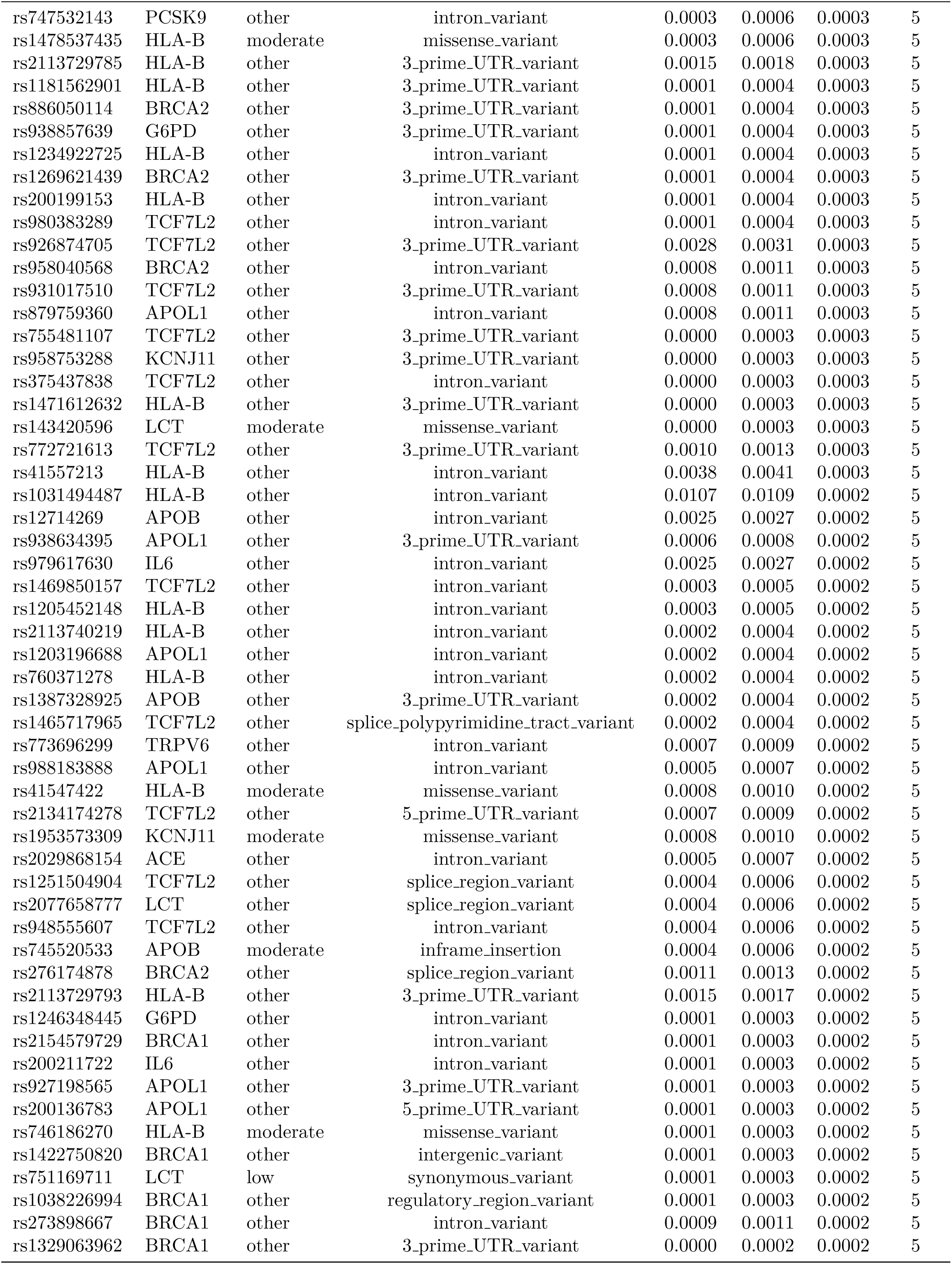
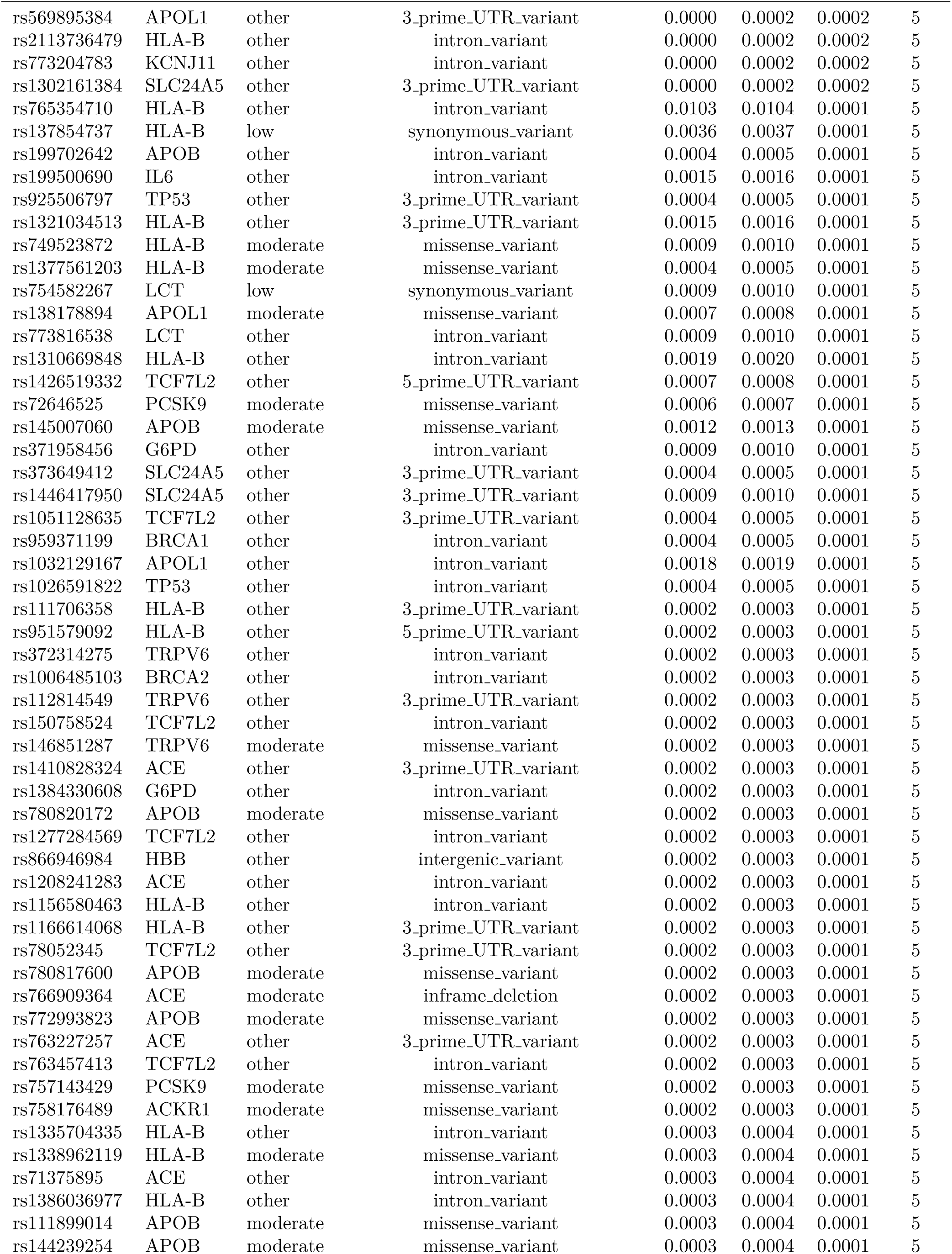
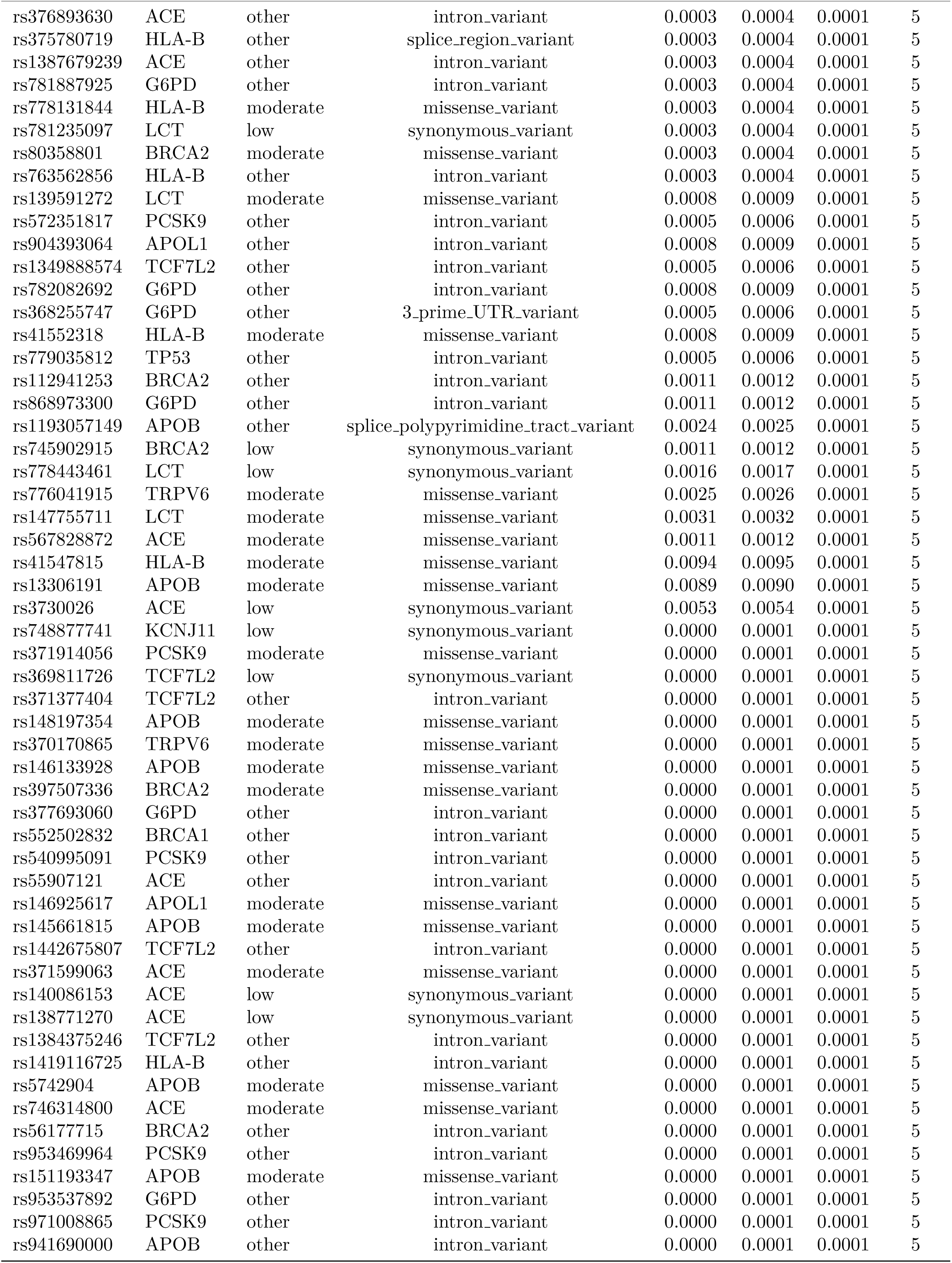
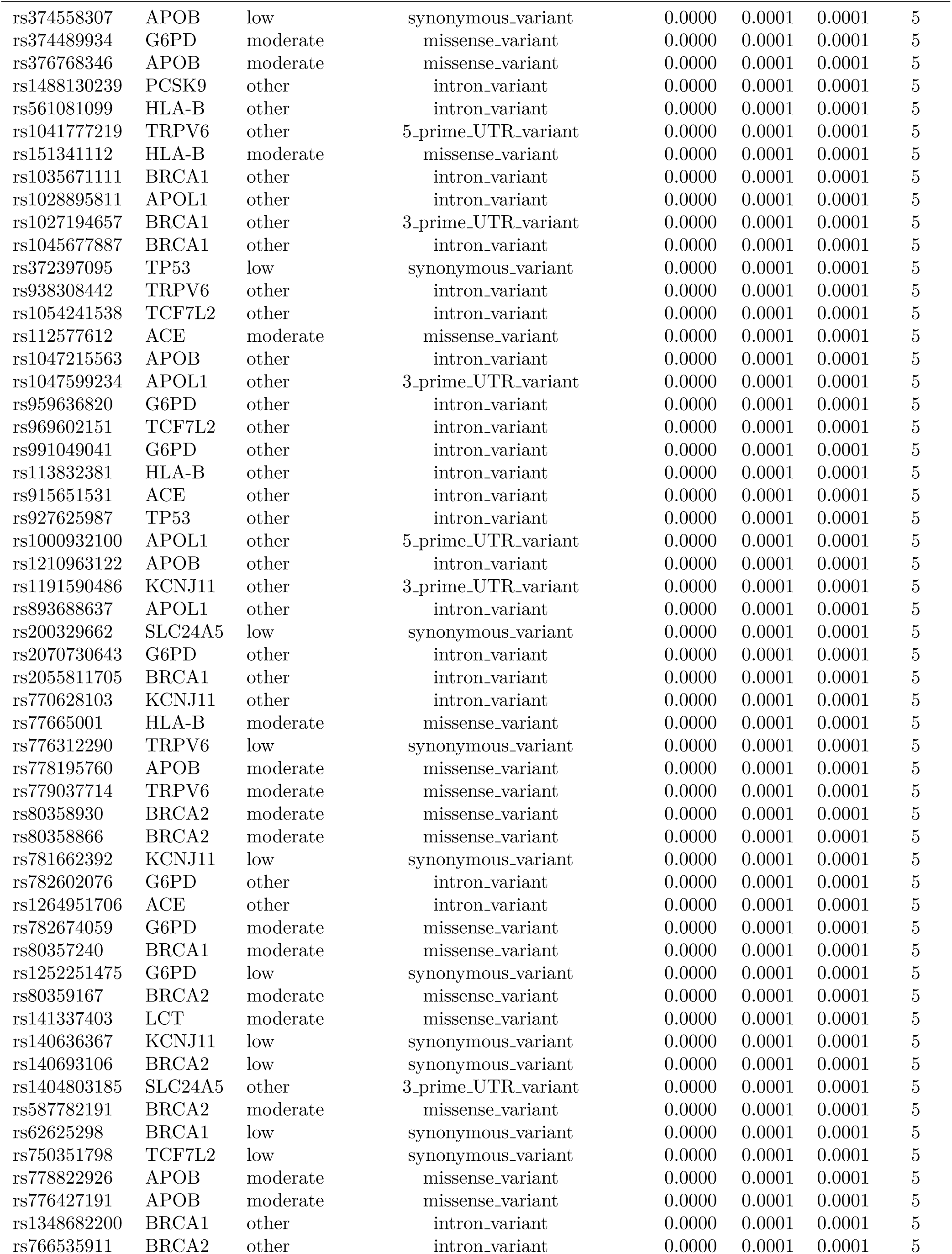
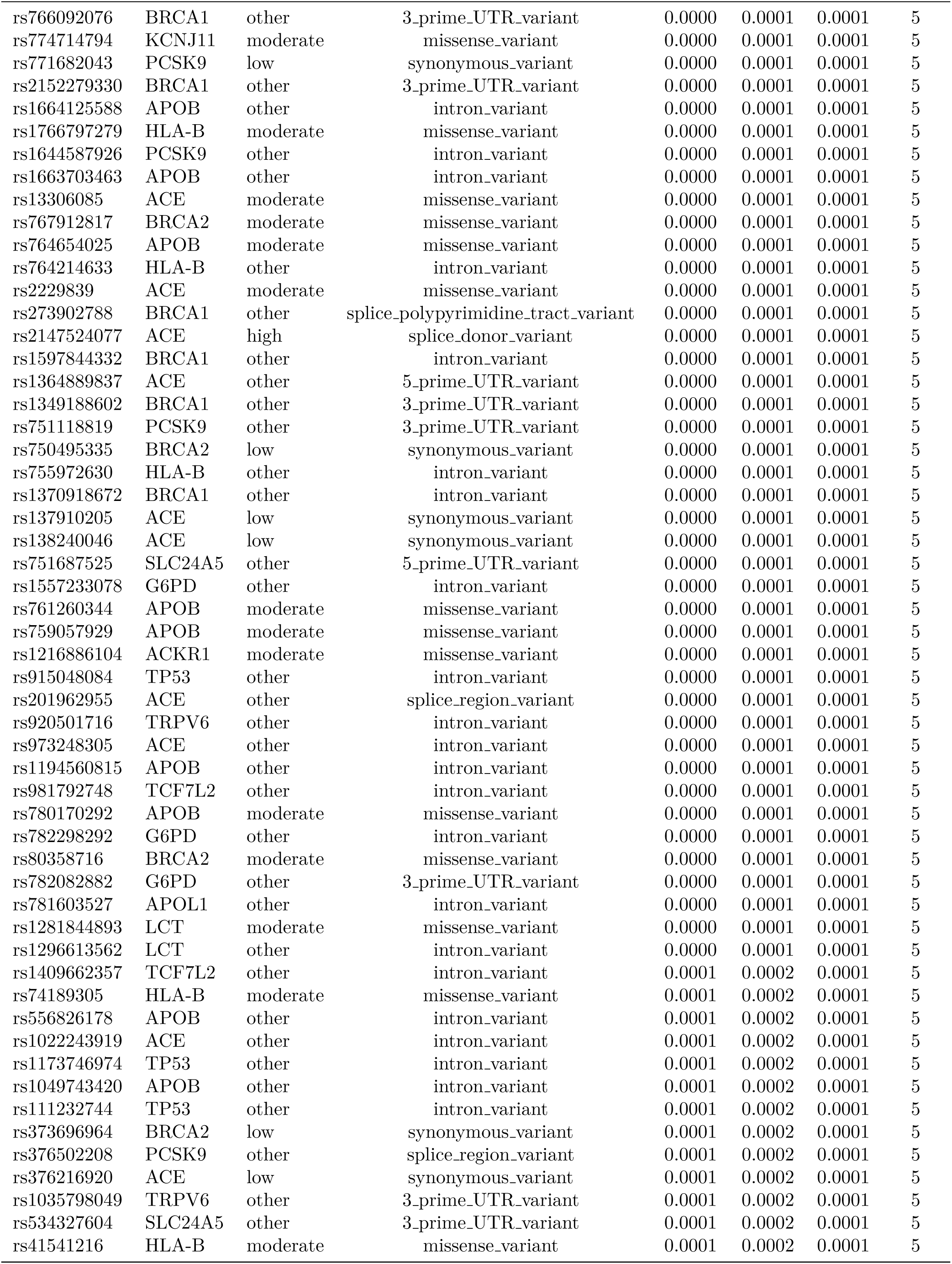
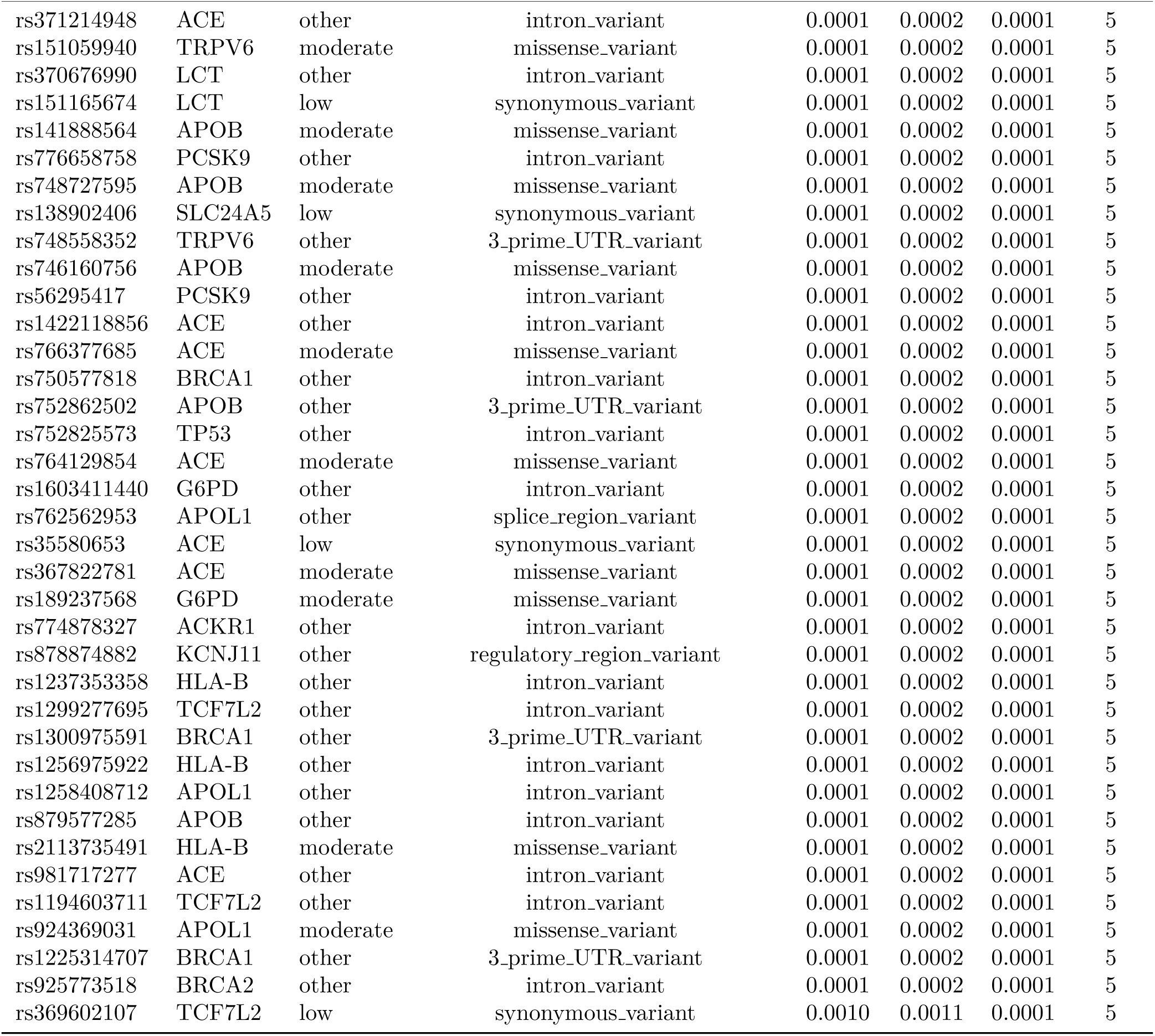
Complete list of methodological disagreements between Naive and Aware estimators.

**Table S3:**
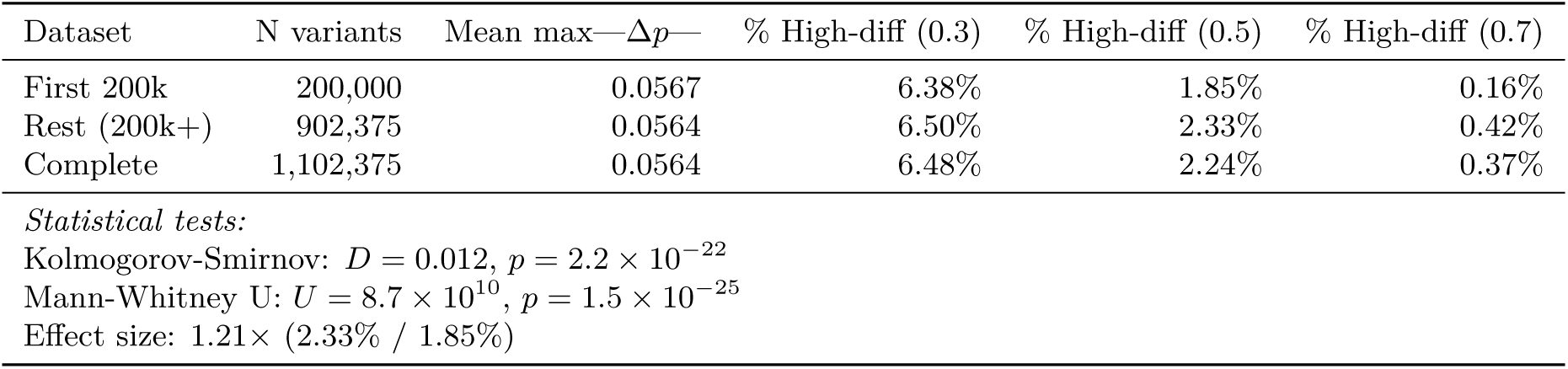
Positional heterogeneity statistics comparing the first 200,000 chromosome 22 variants to the complete dataset.

## Supplementary Methods

### Bootstrap Confidence Interval Methodology

Confidence intervals for odds ratios at the primary differentiation threshold (max—Δ*p*— ≥ 0.5) were estimated using parametric bootstrap resampling. For each of 10,000 iterations, variant counts were resampled from binomial distributions:

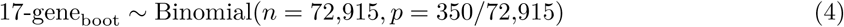

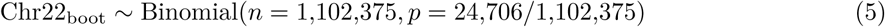

For each resampled dataset, the odds ratio was computed:

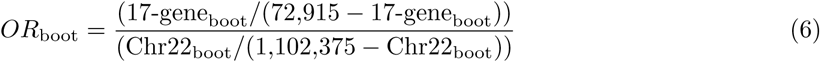

The 95% confidence interval was determined as the 2.5th and 97.5th percentiles of the bootstrap distribution. This approach accounts for sampling uncertainty in both datasets while preserving the observed proportions. The bootstrap distribution was approximately normal (Shapiro-Wilk test *p* = 0.82), validating the parametric approach.

### Chromosome 22 VCF Processing Pipeline

Complete chromosome 22 variant frequencies were computed from 1000 Genomes Project Phase 3 VCF files using cyvcf2 (version 0.30.28). For each variant, genotypes were extracted for all 2,504 samples across 26 populations. For multiallelic variants, alternative allele frequencies were summed per population, and reference allele frequency was computed as:

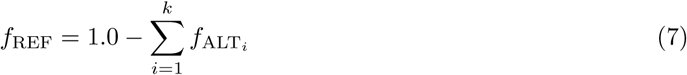

where *k* is the number of alternative alleles. This approach ensures frequencies remain bounded in [0,1] and properly accounts for variants with multiple alternative alleles. Variants with implausible frequencies (*f*_REF_ < −0.01 or *f*_REF_ > 1.01) were excluded (0.1% of raw variants), and remaining frequencies were clipped to [0,1]. The pipeline processed 1,103,547 variants from the source VCF, yielding 1,102,375 variants (99.9% capture) after quality control.

During initial analysis development, a subset of 200,000 variants was processed for computational efficiency. Complete reanalysis revealed only modest positional heterogeneity (Kolmogorov-Smirnov test *p <* 0.001, effect size 1.21×), validating that conclusions were robust to genomic sampling strategy while demonstrating the importance of comprehensive analysis for establishing baseline differentiation rates.

### Reference Allele Frequency Justification

Population differentiation was computed using reference allele frequencies rather than alternative allele frequencies for three reasons. First, each variant has exactly one reference allele (defined by the GRCh38 reference genome), eliminating ambiguity for multiallelic variants where multiple alternative alleles may exist. Second, reference allele frequency is mathematically equivalent to alternative allele frequency for biallelic variants (*f*_REF_+*f*_ALT_ = 1.0), making max—Δ*p*— identical regardless of which allele is measured. Third, reference frequency provides a deterministic, bounded metric that requires no algorithmic choice of which alternative allele to prioritize when multiple alternatives are present.

A sensitivity analysis using a target-alternative-allele approach (selecting the alternative allele with highest mean frequency across populations) yielded 97% concordance in variant-level max—Δ*p*— values (Spearman *ρ* = 0.97, *p <* 10^−300^) and 91.6% agreement in identifying high-differentiation variants (max—Δ*p*— ≥ 0.5), confirming that biological conclusions are robust to allele choice. The 8.4% discordance occurred exclusively among multiallelic variants where alternative alleles showed differential population distributions, with reference-based and alternative-based approaches identifying complementary sets of differentiated loci.

## Supplementary Results

### Coverage Patterns by Gene

Table 1 shows substantial heterogeneity in population coverage across the 17-gene panel. Genes with documented African-population associations showed systematically lower coverage completeness: ACKR1 (7.6% complete), G6PD (4.2%), and HBB (18.5%) compared to genes without strong ancestry-specific associations such as TCF7L2 (61.7%). This pattern likely reflects ascertainment bias in variant discovery, where variants specific to African populations are underrepresented in genotyping arrays designed using European reference panels.

Mean population coverage was 6.39 populations (median 5), with 20,785 variants (28.5%) showing complete 10-population coverage. The bimodal distribution (Figure 1, panel A) suggests two variant classes: (1) common variants discovered in multiple populations and (2) population-specific variants discovered in African samples but not genotyped globally. This ascertainment pattern has implications for interpretation, as the 71.5% of variants with incomplete coverage may include genuinely rare alleles as well as common African-specific alleles absent from non-African panels.

### Correlation Stratified by Coverage Level

When stratified by population coverage, method correlation increased with coverage completeness (Table 3). Low-coverage variants (2-4 populations, *n* = 1,040) showed only 1 disagreement (0.096%), though correlation could not be computed due to insufficient variation. Medium-coverage variants (5-7 populations, *n* = 51,086) showed 1.004% disagreement with strong correlation (*ρ* = 0.9875). High-coverage variants (8-10 populations, *n* = 20,789) showed only 0.019% disagreement (4 variants) with perfect correlation (*ρ* = 1.0000).

This pattern confirms theoretical predictions that method convergence occurs as coverage approaches completeness. The 4 disagreements among high-coverage variants (8-10 populations) all involved variants with exactly 8 or 9 populations, with missing populations contributing extreme frequency differences that naive imputation failed to capture. No variants with complete 10-population coverage showed disagreement, validating that comprehensive population representation eliminates methodological artifacts.

### Gene-Level Enrichment Patterns

Among the 350 high-differentiation variants, enrichment was concentrated in specific genes: TCF7L2 (129 variants, 36.9%), BRCA1 (49, 14.0%), TRPV6 (38, 10.9%), LCT (32, 9.1%), and PCSK9 (29, 8.3%).

This uneven distribution suggests heterogeneity in selection pressures across the panel. TCF7L2’s disproportionate representation likely reflects the gene’s large size (215 kb, 10,258 variants in dataset) combined with extensive linkage disequilibrium, where selection on a single causal variant sweeps linked neutral variants to high frequency. LCT enrichment aligns with documented selection for lactase persistence in pastoralist populations. The concentration in specific genes supports the interpretation that high differentiation reflects localized positive selection rather than relaxed constraint across entire genes.

## Supplementary Data Files

The following data files are available for download https://osf.io/7vgcy/files/osfstorage/69540811ebe8f83cffa69

- **tableS2 all disagreements.csv**: Complete list of 518 variants showing disagreement between methods, including variant IDs, gene names, differentiation scores, delta values, and population coverage.
- **chr22 complete metrics FINAL.csv**: Complete differentiation metrics for all 1,102,375 chromosome 22 variants, including variant positions, reference alleles, naive and missingness-aware max—Δ*p*— values, and population coverage.
- **enrichment analysis.csv**: Enrichment statistics across five differentiation thresholds (0.3, 0.4, 0.5, 0.6, 0.7), including odds ratios, p-values, and fold-change interpretations.
- **bootstrap ci.csv**: Bootstrap confidence interval estimates for the primary threshold (0.5), including observed OR, bootstrap mean, bootstrap median, and 95% CI bounds.

